# A theory of multineuronal dimensionality, dynamics and measurement

**DOI:** 10.1101/214262

**Authors:** Peiran Gao, Eric Trautmann, Byron Yu, Gopal Santhanam, Stephen Ryu, Krishna Shenoy, Surya Ganguli

## Abstract

In many experiments, neuroscientists tightly control behavior, record many trials, and obtain trial-averaged firing rates from hundreds of neurons in circuits containing billions of behaviorally relevant neurons. Dimensionality reduction methods reveal a striking simplicity underlying such multi-neuronal data: they can be reduced to a low-dimensional space, and the resulting neural trajectories in this space yield a remarkably insightful dynamical portrait of circuit computation. This simplicity raises profound and timely conceptual questions. What are its origins and its implications for the complexity of neural dynamics? How would the situation change if we recorded more neurons? When, if at all, can we trust dynamical portraits obtained from measuring an infinitesimal fraction of task relevant neurons? We present a theory that answers these questions, and test it using physiological recordings from reaching monkeys. This theory reveals conceptual insights into how task complexity governs both neural dimensionality and accurate recovery of dynamic portraits, thereby providing quantitative guidelines for future large-scale experimental design.

## 1 Introduction

In this work, we aim to address a major conceptual elephant residing within almost all studies in modern systems neurophysiology. Namely, how can we record on the order of hundreds of neurons in regions deep within the brain, far from the sensory and motor peripheries, like mammalian hippocampus, or pre-frontal, parietal, or motor cortices, and obtain scientifically interpretable results that relate neural activity to behavior and cognition? Our apparent success at this endeavor seems absolutely remarkable, considering such circuits mediating complex sensory, motor and cognitive behaviors contain *O*(10^6^) to *O*(10^9^) neurons [Shepherd, 2004] - 4 to 7 orders of magnitude more than we currently record. Or alternatively, we could be completely misleading ourselves: perhaps we should not trust scientific conclusions drawn from statistical analyses of so few neurons, as such conclusions might become qualitatively different as we record more. Without an adequate theory of neural measurement, it is impossible to *quantitatively* adjudicate where systems neuroscience currently stands between these two extreme scenarios of success and failure.

One potential solution is an experimental one: simply wait until we can record more neurons. Indeed, exciting advances in recording technology over the last several decades have lead to a type of Moore’s law in neuroscience: an exponential growth in the number of neurons we can simultaneously record with a doubling rate of 7.4 years since the 1960’s [Stevenson and Kording, 2011]. Important efforts like the BRAIN Initiative promise to ensure such growth in the future. However, if we simply extrapolate the doubling rate achieved over the last 50 years, we will require about 100 to 200 years to record 4 to 7 orders of magnitude more neurons. Thus, for the near future, it is highly likely that measurements of neural dynamics at single cell, single spike-time resolution in mammalian circuits controlling complex behaviors will remain in the highly sub-sampled measurement regime. Therefore we need a theory of neural measurement that addresses a fundamental question: how and when do statistical analyses applied to an infinitesimally small subset of neurons reflect the collective dynamics of the much larger, unobserved circuit they are embedded in?

Here we provide the beginnings of such a theory, that is quantitatively powerful enough to (a) formulate this question with mathematical precision, (b) make well defined, testable predictions that guide the interpretation of past experiments, and (c) provide a theoretical framework to guide the design of future large scale recording experiments. We focus in this work on an extremely commonly used experimental design in which neuroscientists repeat a given behavioral or cognitive task over many trials, and record the trial averaged neural dynamics of many neurons. An advantage of this design, which has promoted its widespread usage, is that the neurons need not be simultaneously recorded. This resulting trial average firing rate dynamics can be thought of as a collection of neural trajectories exploring a high dimensional neural space, with dimensionality equal to the number of recorded neurons (see e.g. Fig 1 below for a conceptual overview). They reflect a fundamental description of the state space dynamics of the neural circuit during cognition and behavior. Almost always, such trajectories are analyzed via dimensionality reduction (see [Cunningham and Yu, 2014] for a review), and almost ubiquitously, a large fraction of variance in these trajectories lives in a much lower dimensional space.

**Figure 1:**
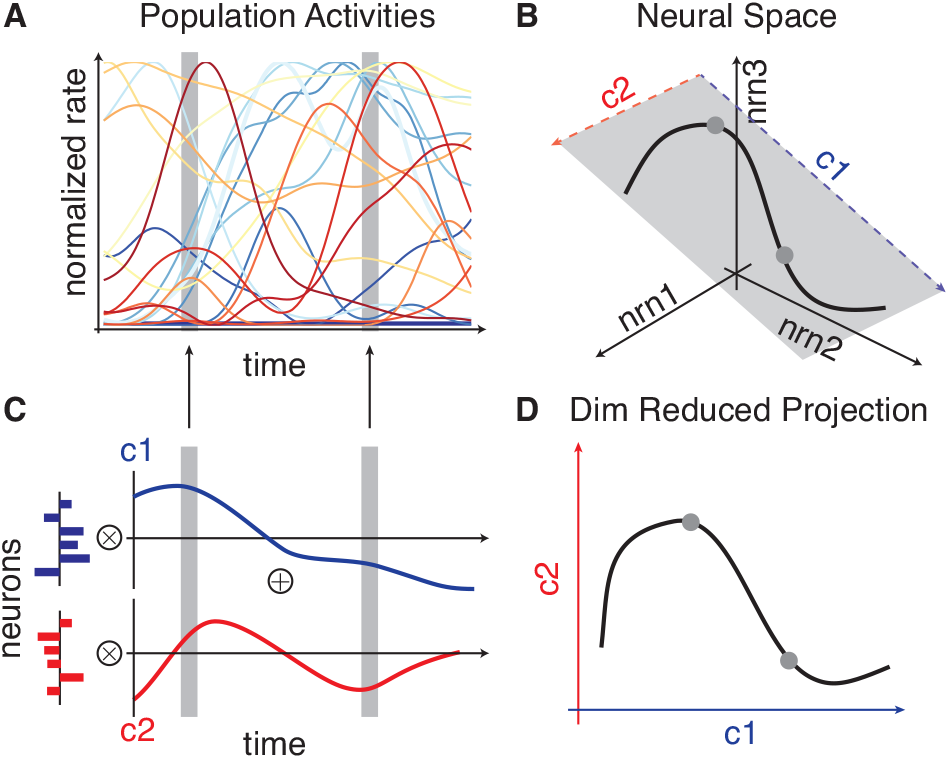
Conceptual overview of state space dynamics through dimensionality reduction. **(A)** The trial averaged firing rate dynamics of *M* neurons. **(B)** This data can be equivalently thought of as tracing out a neural state space trajectory in an *M* dimensional firing rate space with one axis per neuron. Here only three of the *M* axes are shown, and as illustrated, sometimes such a trajectory can be largely confined to a lower dimensional subspace, here a two dimensional subspace. **(C)** A decomposition of the data in **(A)** into two static spatial patterns across neurons (red and blue patterns, left). The population activity pattern at each instant of time is a weighted sum of the two static *basis* patterns, where the weighting coefficients (red and blue traces, right) depend on time. If all population patterns across time can largely be explained by linear combinations of these two patterns, then the neural trajectory corresponding to the data in **(A)** will be largely confined to explore a two dimensional subspace within *M* dimensional firing rate space, shown for example in **B**. Two special points in the subspace correspond to the two basis patterns in **(C)**, i.e. the endpoints of the two red and blue vectors in **(B)**, and the subspace is spanned by all linear combinations of these two patterns. **(D)**. One can then accurately visualize the state space dynamics in pattern space, by plotting the time-dependent weighting coefficients in **(C)** against each other. Each axis in pattern space corresponds to how much each basis pattern is present in the neural population.

The resulting neural trajectories in the low dimensional space often provide a remarkably insightful dynamical portrait of circuit computation during the task in a way that is inaccessible through the analysis of individual neurons [Briggman et al., 2006]. For example, curvature in the geometry of these dynamical portraits recovered from macaque prefrontal cortex by [Mante et al., 2013] revealed a novel computational mechanism for contextually-dependent integration of sensory evidence. Similarly, dimensionality reduction by [Machens et al., 2010] uncovered dynamical portraits which revealed that macaque somatosensory cortices compute both stimulus frequency and time in a functionally but not anatomically separable manner in a tactile discrimination task. Dynamical portraits obtained by [Mazor and Laurent, 2005] revealed that neural transients in insect olfactory areas rapidly computed odor identity long before the circuit settled to a steady state. And an analysis of neural dynamics in macaque parietal cortex showed that the dynamical portraits were largely one-dimensional, revealing an emergent circuit mechanism for robust timing in attentional switching and decision making [Ganguli et al., 2008a]. Also, the low-dimensional activity patterns found in primary motor-cortex provide causal constraints learning itself [Sadtler et al., 2014].

Given the importance of these dynamical portraits as a first window into circuit computation, it is important to ask if we can trust them despite recording so few neurons? For example, would their geometry change if we record more neurons? How about their dimensionality? The ubiquitous low dimensionality of neural recordings suggests an underlying simplicity to neural dynamics; what is its origin? How does the number of neurons we need to record to accurately recover dynamical portraits scale with the complexity of the task, and properties of the neural dynamics? Indeed which minimal properties of neural dynamics are important to know in order to formulate and answer this last question?

Our theory provides a complete answer to these questions within the context of trial averaged experimental design. Central to our theory are two main conceptual advances. The first is the introduction of neural task complexity (NTC), a mathematically well defined quantity that takes into account both the complexity of the task, and the smoothness of neural trajectories across task parameters. Intuitively, the NTC measures the volume of the manifold of task parameters, in units of the length scales over which neural trajectories vary across task parameters, and it will be small if tasks are very simple and neural trajectories are very smooth. We prove that this measure upper bounds the dimensionality of neural state space dynamics. This theorem has important implications for systems neuroscience: it is likely that the ubiquitous low dimensionality of measured neural state space dynamics is due to a small NTC. In any such scenario, simply recording many more neurons than the NTC, while repeating the same simple task will not lead to richer, higher dimensional datasets; indeed data dimensionality will become independent of the number of recorded neurons. One would have to move to more complex tasks to obtain more complex, higher dimensional dynamical portraits of circuit computation.

The second conceptual advance is a novel theoretical link between the act of neural measurement and a technique for dimensionality reduction known as random projections. This link allows us to prove that, as long as neural trajectories are sufficiently randomly oriented in state space, we need only record a number of neurons proportional to the product of the number of task parameters and the *logarithm* of the NTC. This theorem again has significant implications for systems neuroscience. Indeed, it quantitatively adjudicates between the two extremes of success or failure raised above, fortunately, in the direction of success: it is highly likely that low dimensional dynamical portraits recovered from past experiments are reasonably accurate despite recording so few neurons, because those tasks were so simple, leading to a small NTC. Moreover, as we begin to move to more complex tasks, this theorem provides rigorous guidance for how many more neurons we will need to record in order to accurately recover the resulting more complex, higher dimensional dynamical portraits of circuit computation.

Below, we build up our theory step by step. We first review the process of recovering state space dynamical portraits through dimensionality reduction in neuroscience. We then introduce the notion of NTC, and illustrate how it provides an upper bound on neural dimensionality. Then we review the notion of random projections, and illustrate how the NTC of an experiment *also* determines how many neurons we must record to accurately obtain dynamical portraits. Along the way, we extract a series of experimentally testable predictions, which we confirm in neural recordings from the motor and premotor cortices of monkeys performing reaches to multiple directions. We end in the discussion with an intuitive summary of our theory and its implications for the future of large scale recordings in systems neuroscience.

## 2 Recovery of neural state space dynamics through dimensionality reduction

Imagine an experiment in which a neuroscientist records trial averaged patterns of neural activity from a set of *M* neurons across time. We denote by **x**_*i*_(*t*) the trial averaged firing rate of neuron *i* at time *t*. These data are often visualized by superimposing the firing rates of each neuron across time (Fig. 1A). Alternatively, these data can be thought of as a neural trajectory in an *M* dimensional space (Fig. 1B). At each time, the measured state of the neural circuit consists of the instantaneous pattern of activity across *M* neurons, which corresponds to a point in *M* dimensional space, where each dimension, or axis in the space corresponds to the firing rate of one neuron. As time progresses, this point moves, tracing out the neural trajectory.

It is difficult to directly understand or visualize this trajectory, as it evolves in such a high-dimensional ambient space. Here dimensionality reduction is often used to simplify the picture. The main idea behind many linear dimensionality reduction methods is to decompose the entire set of dynamic neural activity patterns across neurons, unfolding over time, into a time dependent linear combination of a fixed set of static patterns across neurons (Fig. 1C). The hope is that the data can be dramatically simplified if a linear combination of a small number of static basis patterns are sufficient to account for a large fraction of variance in the data across time. When this is the case, the neural state space dynamics can unfold over time in a much lower dimensional pattern space, whose coordinates consist of how much each static pattern is present in the neural population (Fig. 1D).

Mathematically, the decomposition illustrated in Fig. 1C can be written as 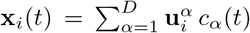, where each *M* dimensional vector **u**^*α*^ is a static basis pattern across neurons, each *c*_*α*_(*t*) is the amplitude of that pattern across time (Fig.1C), and *D* denotes the number of patterns or dimensions retained. Principal components analysis (PCA) is a simple way to obtain such a decomposition (see Supplementary Material for a review). PCA yields a sequence of basis patterns **u**^*α*^, *α* = 1, …, *M* each accounting for a different amount of variance *µ*^*α*^ in the neural population. The patterns can be ordered in decreasing amount of variance explained, so that *µ*_1_ ≥ *µ*_2_ ≥, …, ≥ *µ*_*M*_. By retaining the top *D* patterns, one achieves a fraction of variance explained given by 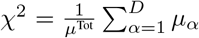, where 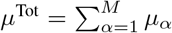 is the total variance in the neural population. Dimensionality reduction is considered successful if a small number of patterns *D* relative to number of recorded neurons *M*, accounts for a large fraction of variance explained in the neural state space dynamics.

How well does dimensionality reduction perform in practice in neurophysiology data? We have performed a meta-analysis (Fig. 2) of a diverse set of 20 experiments spanning a variety of model organisms (macaques, rodents, and insects), brain regions (hippocampal, prefrontal, parietal, somatosensory, motor, premotor, visual, olfactory and brainstem areas), and tasks (memory, decision making, sensory detection and motor control). This meta-analysis reveals that dimensionality reduction as a method for simplifying neural population recordings performs exceedingly well. Indeed it reflects one of the most salient aspects of systems neurophysiology to have emerged over the last couple of decades: namely that neuronal recordings are often far lower dimensional than the number of recorded neurons. Moreover, in each of these works, the associated low dimensional dynamical portraits provide insights into relations between neural population activity and behavior. Despite this almost ubiquitous simplicity found in neural population recordings, prior to this work, we are unaware of any simple, experimentally testable theory that can quantitatively explain the dimensionality and accuracy of these recovered dynamical portraits.

**Figure 2:**
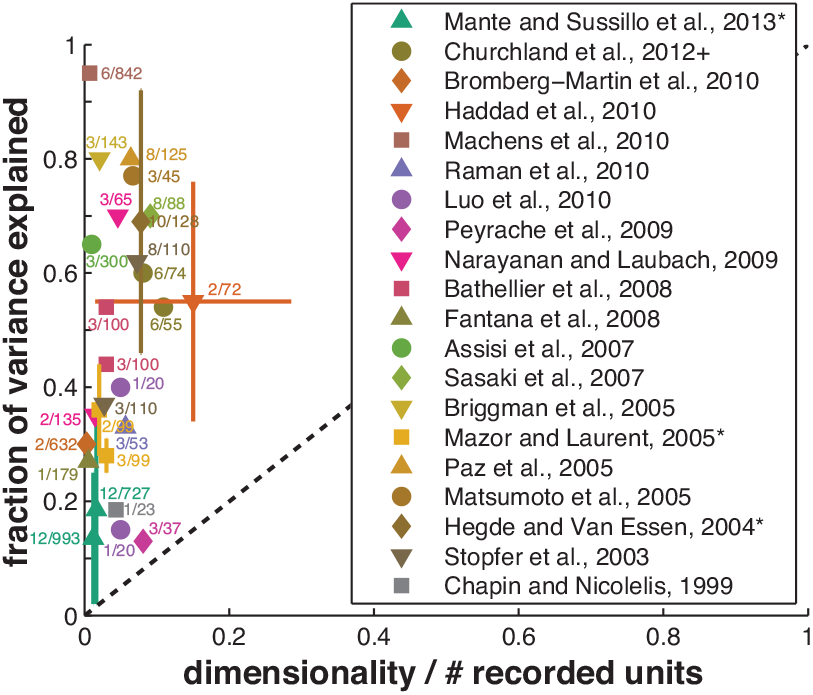
The ubiquity of low dimensionality relative to number of recorded neurons. In many experiments (e.g. in insect [Stopfer et al., 2003, Mazor and Laurent, 2005, Assisi et al., 2007, Raman et al., 2010, Haddad et al., 2010] olfactory systems, mammalian olfactory [Bathellier et al., 2008, Haddad et al., 2010], pre-frontal [Narayanan and Laubach, 2009, Peyrache et al., 2009, Machens et al., 2010, Warden and Miller, 2010, Mante et al., 2013], motor and premotor,[Paz et al., 2005, Churchland et al., 2012], somatosensory [Chapin and Nicolelis, 1999], visual [Hegde´ and Van Essen, 2004, Matsumoto et al., 2005], hippocampal [Sasaki et al., 2007], and brain stem [Bromberg-Martin et al., 2010] systems) a *much* smaller number of dimensions than the number of recorded neurons captures a large amount of variance in neural firing rates.

## 3 Neural Task Complexity and Dimensionality

We now begin to describe such a theory. Central to our theory is the notion of neural task complexity (NTC), which both upper bounds the dimensionality of state space dynamics and quantifies how many neurons are required to accurately recover this dynamics. Here, we first consider the dimensionality of the dynamics, and later we consider the accuracy of the dynamics. To introduce the NTC intuitively, imagine how many dimensions a single neural trajectory could possibly explore. Consider for concreteness, the trial averaged neural population activity while a monkey is performing a simple reach to a target (Fig. 3AB). This average reach lasts a finite amount of time *T*, which for example could be about 600ms. The corresponding neural trajectory (Fig. 3C) can explore neural space for this much time, but it cannot change direction infinitely fast. The population response is limited by an autocorrelation time *τ* (see supplementary methods for details). Roughly, one has to wait an amount of time *τ* before the neural population’s activity pattern changes appreciably (Fig. 3B) and therefore the neural trajectory can bend to explore another dimension (Fig. 3C). This implies that the maximal number of dimensions the state space dynamics can possibly explore is proportional to *T/τ*. Of course the constant of proportionality is crucial, and our theory, applicable to reaching data described below, computes this constant to be 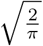 (see supplementary material for a proof and a definition of *τ*), yielding, for this experiment, an 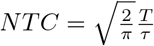.

**Figure 3:**
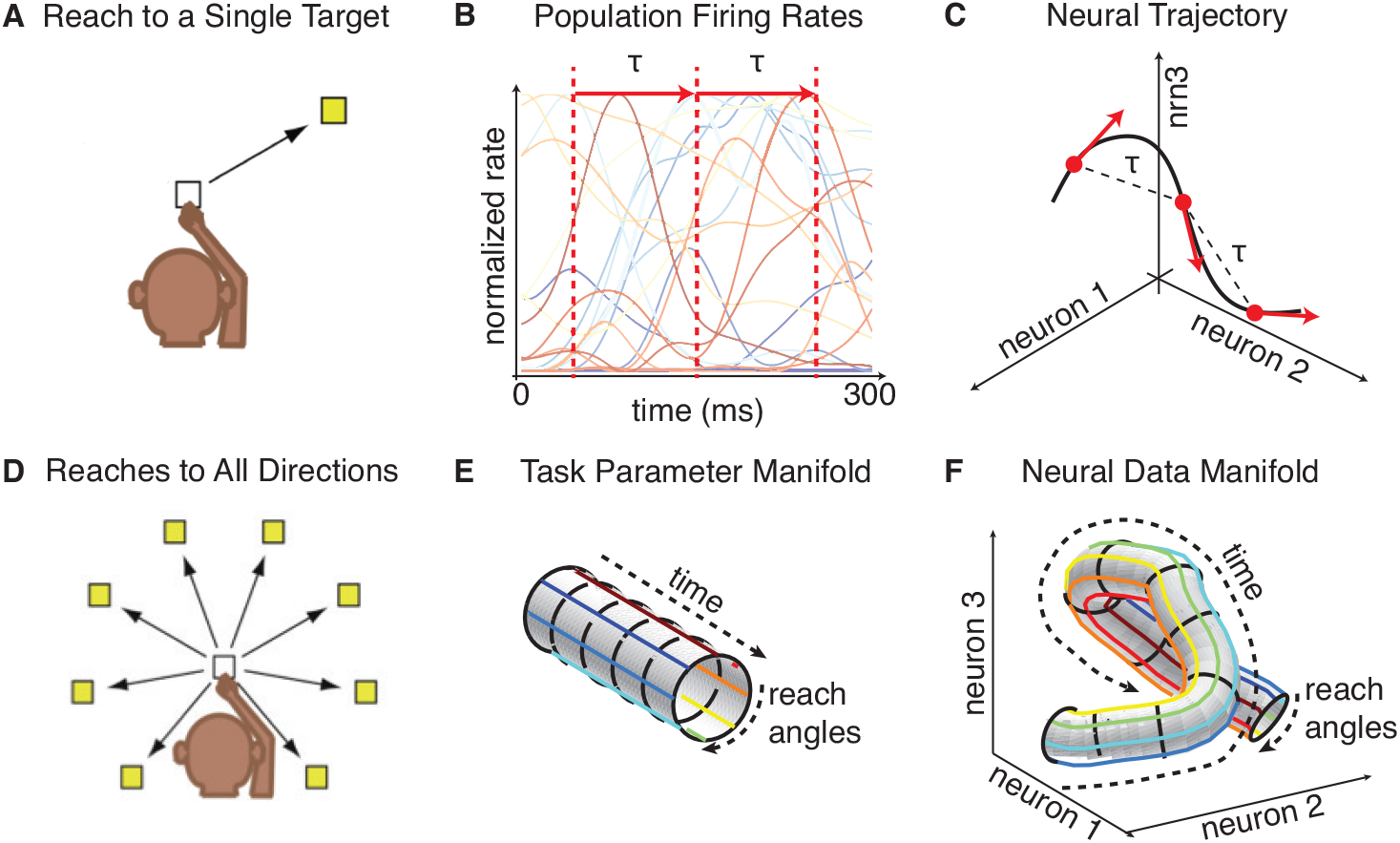
Trial averaged neurophysiology as an embedding of a task manifold into a neural manifold. **(A)** A monkey reaching to a single target. The task is parameterized simply by time *t*. **(B)** Schematic of neuronal firing rate data during the reach, and **(C)** the associated neural trajectory. **(D)** A monkey reaching to several targets. **(E)** The associated task manifold is a cylinder parameterized by time into the reach and reach angle. **(F)** The neural state space dynamics is a smooth embedding of the task manifold into neural firing rate space.

Now most tasks have more than just time as a parameter. Consider a slightly more complex experiment in which a monkey reaches to 8 different targets (Fig. 3D). Now the manifold of trial averaged task parameters is a cylinder, parameterized by time *t* into the reach and reach angle *θ* (Fig. 3E). Since for each time *t* and angle *θ*, there is a corresponding trial averaged neural activity pattern across neurons **x**_*i*_(*θ, t*), the neural state space dynamics is fundamentally an embedding of this task manifold into neural space, yielding a curved intrinsically two dimensional neural manifold that could potentially explore many more than two dimensions in firing rate space by curving in different ways (Fig. 3F). How many dimensions could it possibly explore? Well the same argument that we made for time into a reach at fixed angle, also holds for reaching across all angles at a fixed time into the reach. The total extent of angle is 2*π*. Moreover, the neural population response cannot vary infinitely fast across angle; it has a spatial autocorrelation length ∆. Intuitively, this means that the two patterns of activity **x**_*i*_(*θ*_1_*, t*) and **x**_*i*_(*θ*_2_*, t*) across neurons at two different reach angles *θ*_1_ and *θ*_2_ at the same time *t* will not be appreciably different unless |*θ*_*i*_ − *θ*_*j*_| > ∆. Roughly, one can think of ∆ as the average width of single neuron tuning curves across reach angle.

Thus, just as in the argument for time, because the total angle to be explored is limited to 2*π*, and patterns are largely similar across angular separations less than ∆, the maximal number of dimensions a single circle around the neural manifold at any fixed time could explore, is proportional to 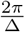, where the proportionality constant is again 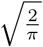. Now intuitively, the number of dimensions the full neural manifold could explore across both time and angle would be maximized if these explorations were independent of each other. Then the maximal dimensionality would be the product of 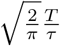 and 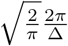 (see supplementary material for a proof), yielding an 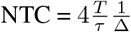.

More generally, consider a task that has *K* task parameters indexed by *k* = 1*, …, K*, each of which vary over a range *L_k_*, and for which neural activity patterns have a correlation length *λ_k_*. Then the NTC is

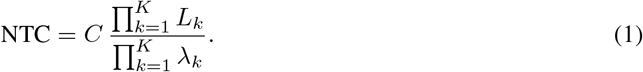

For example, in the special case of reaches to all angles we have considered so far, we have *K* = 2, *L*_1_ = *T*, *λ*_1_ = *τ*, *L*_2_ = 2*π*, *λ*_2_ = ∆, and 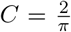. In the general case, our theory (see supplementary material) pro-vides a precise method to define the autocorrelation lengths *λ*_*k*_, in a manner consistent with the intuition that a correlation length measures how far one has to move in behavioral manifold, to obtain an appreciably different pattern of activity across neurons in the neural manifold (Fig. 3EF). Moreover, our theory determines the constant of proportionality *C*, as well as provides a proof that the neural dimensionality *D*, measured by the participation ratio of the PCA eigenvalue spectrum (see methods) is less than the minimum of the number of recorded neurons *M* and the *NTC*:

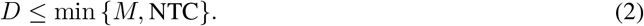

Also, in the supplementary material, we consider a much simpler scenario in which there are a finite set of *P* neural activity patterns, for example in response to a finite set of *P* stimuli. There, the NTC is simply *P*, and we compute analytically how measured dimensionality *D* increases with number of recorded neurons *M*, and how it eventually asymptotes to the NTC if there are no further constraints on the neural representation. In the following, however, we focus on the much more interesting case of neural manifolds in (1).

We note that the NTC in (1) takes into account two very distinct pieces of information. First, the numerator only knows about the task design; indeed it is simply the volume of the manifold of task parameters (e.g. Fig. 3E). The denominator on the other hand requires knowledge of the smoothness of the neural manifold; indeed it is the volume of an autocorrelation cell over which population neural activity does not change appreciably across *all K* task parameters. Thus the theorem (2) captures the simple intuition that neural manifolds of limited volume and curvature (e.g. Fig. 3CF) cannot explore that many dimensions (though they could definitely explore fewer than the NTC). However, as we see below the precise theorem goes far beyond this simple intuition, as it provides a quantitative framework to guide the interpretation of past experiments and design future ones.

## 4 A Dimensionality Frontier in Motor Cortical Data

To illustrate the interpretative power of the NTC, we re-examined the measured dimensionality of neural activity from the motor and premotor cortices of two monkeys, H and G, recorded in [Yu et al., 2007], during an eight-direction center-out reach task, as in Fig. 3D (see also, Methods). The dimensionality of the entire dataset, i.e. the number of dimensions explored by the neural manifold in Fig. 3F, was 7.1 for monkey H and 4.6 for monkey G. This number is far less than the number of recorded single units, which were 109 and 42 for monkeys H and G respectively. So a natural question is, how can we explain this order of magnitude discrepancy between the number of recorded units and the neural dimensionality, and would the dimensionality at least increase if we recorded more units? In essence, what is the origin of the simplicity implied by the low dimensionality of the neural recordings?

Here, our theorem (2) can provide conceptual guidance. As illustrated in Fig. 4A, our theorem in general implies that experiments in systems neurophysiology can live within 3 qualitatively distinct experimental regimes, each with its own unique predictions. First, the dimensionality *D* could be close to the number of recorded neurons *M* but far less than the NTC. This scenario suggests one may not be recording enough neurons, and that the dimensionality and accuracy of dynamic portraits may increase when recording more neurons. Second, the dimensionality may be close to the NTC but far below the number of neurons. This suggests that the task is very simple, and that the neural dynamics is very smooth. Recording more neurons would not lead to richer, higher dimensional trial averaged dynamics; the only way to obtain richer dynamics, at least as measured by dimensionality, is to move to a more complex task. Finally, and perhaps most interestingly, in the third regime, dimensionality may be far less than both the NTC and the number of recorded neurons. Then, and only then, can one say that the dimensionality of neural state space dynamics is constrained by neural circuit properties above and beyond constraints imposed by the simplicity of the task and the smoothness of the dynamics alone.

**Figure 4:**
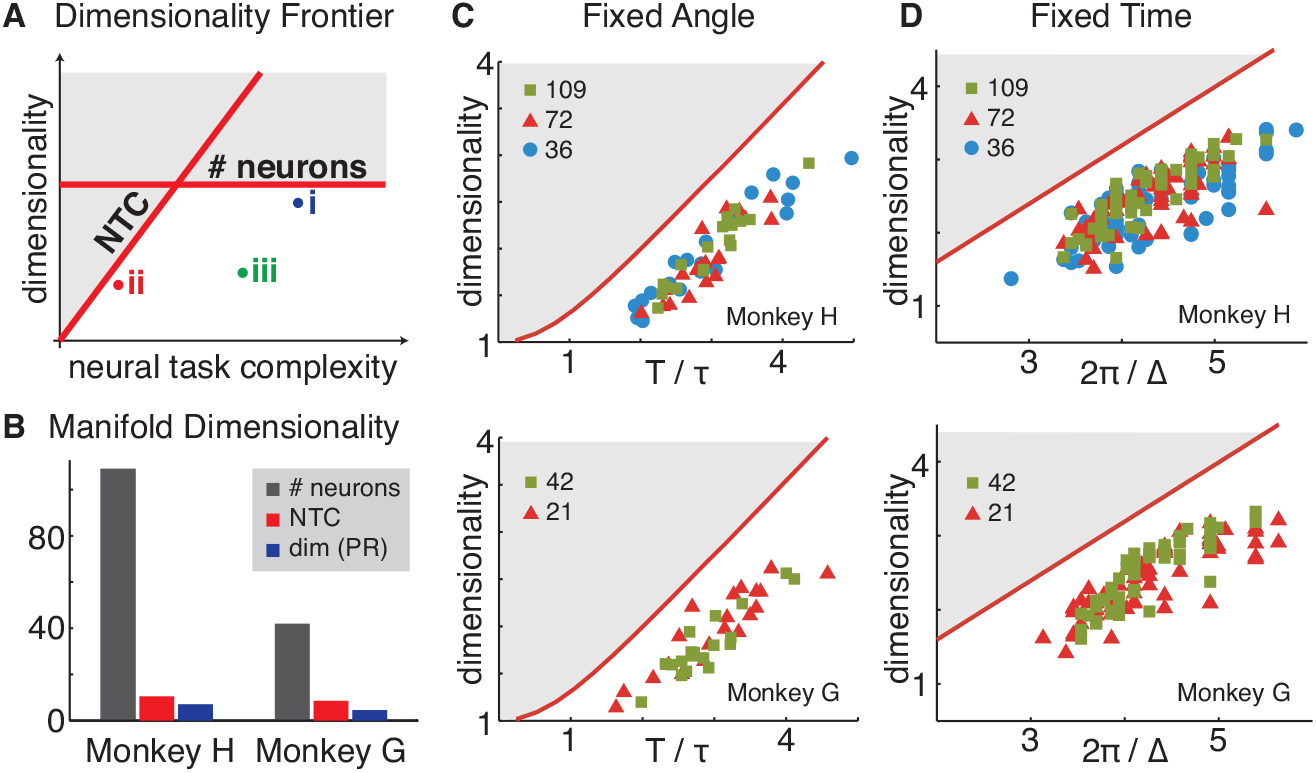
A Dimensionality frontier in motor cortical data. **(A)** A schematic of the dimensionality frontier imposed by theorem (2). Allowed possibilities of dimensionality *D* and neural task complexity, NTC, exhibit 3 distinct regimes: (i) the number of recorded neurons *M* but not NTC restricts dimensionality, (ii) NTC but not *M* restricts *D*, and (iii) *D*is far less than both *M* and *NTC*, reflecting an unexplained circuit constraint beyond smoothness and task simplicity. **(B)** The number of recorded neurons *M*, neural task complexity, NTC and dimensionality *D* are 109, 10.5, and 7.1 respectively for Monkey H, and 42, 8.6 and 4.6 for monkey G. **(C)** Dimensionality and NTC as we explore segments of neural trajectories of different durations at fixed angle, on the full neural manifold in Fig 3E. Temporal segments of different durations were chosen from a uniform distribution between 100ms and 600ms from all reach angles. For each segment we computed its dimensionality and plotted it against its own 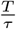 ratio to obtain the scatter plot. The red curve indicates the dimensionality frontier, predicted by theory to be proportional to 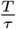. **(D)** Dimensionality and NTC as we explore along all angles on the neural manifold in 3E at fixed times. Relative to movement onset, at each 10ms interval we computed the number of dimensions explored across angles against 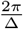 to obtain the scatter plot. Due to the natural variability in the time-dependent population tuning width, we were able to obtain ∆’s of different values. The red curve indicates the dimensionality frontier, predicted by theory to be proportional to 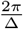. Moreover in both **(C)** and **(D)**, we repeat the above analyses, while discarding a randomly chosen one third (red triangles) or two thirds (blue circles) of the recorded neurons for monkey H (top panels) and one half (red triangles) of the recorded neurons for monkey G (bottom panels).

Returning to the motor cortical data, it is clear that scenario (i) is ruled out in this experiment. But without the definition and computation of the NTC, one cannot distinguish between scenarios (ii) and (iii). We computed the spatial and temporal autocorrelation lengths of neural activity across reach angle and time, and found them to be ∆ = 1.82 radians and *τ* = 126 ms in monkey H, and ∆ = 1.91 radians and *τ* = 146 ms in monkey G. Given that the average reach duration is *T* = 600 ms in both monkeys, the 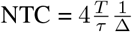 is 10.5 for monkey H and 8.6 for monkey G. Comparing these NTCs to the dimensionalities *D* = 7.1 for monkey H and *D* = 4.6 for monkey G, and the number of recorded neurons *M* = 109 for monkey H and *M* = 42 for monkey G (see Fig. 4B), we see that this experiment is deep within experimental regime (ii) in Fig. 4A.

This deduction implies several striking predictions for motor cortical dynamics. First, assuming the rest of the unrecorded neurons are statistically homogenous to the recorded neurons (implying that population smoothness *τ* and ∆ would not change much as we added more neurons to our measured population), then if we were to record more neurons, even roughly all 500 million neurons in macaque motor cortex, the dimensionality of the neural manifold in each monkey would not exceed 10.5 and 8.6 respectively.

Equivalently, if we were to drop a significant fraction of neurons from the population, the dimensionality would remain essentially the same. In essence, dimensionality would be largely *independent* of the number of recorded neurons. The second prediction is that if we were to vary the NTC, by varying the task, then this would have a significant impact on dimensionality: it would be proportional to the NTC.

We confirm both of these predictions in Fig. 4CD. First, in the given dataset, we cannot increase the NTC further, but we can reduce it by restricting our attention to subsets of reach extents and angles. In essence we explore restricted one-dimensional slices of the full neural manifold in Fig. 3F as follows. First, in Fig. 4B, we explore different random time intervals at different fixed angles, and we plot the dimensionality explored by the segment of neural trajectory against the duration *T* of the trajectory divided by its autocorrelation *τ*. Moreover, we vary the number of recorded neurons we keep in our analysis. Second, in Fig. 4C, we pick different times and we plot the number of dimensions explored by the neural manifold (now a circle) across all angles at each chosen time, against 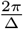, where ∆ is the smoothness parameter of the neural circle at that time, again also varying the number of neurons in our analysis. As can be clearly seen, in both monkeys the predictions of the theory in experimental regime (ii) in Fig. 4A are confirmed: dimensionality is largely independent of the number of recorded neurons, and it hugs closely the dimensionality frontier set by the NTC.

Overall, these results suggest a conceptual revision in the way we may wish to think about neural complexity as measured by its dimensionality. Essentially, neural state space dynamics should not be thought of as inherently simple just because its measured dimensionality is much less than the number of recorded neurons. Instead, by properly comparing dimensionality to neural task complexity, we find neural state space dynamics in motor cortex is as complex and as high-dimensional as possible given basic task constraints and neural smoothness constraints. In essence, the neural state space dynamics represented in Fig. 3F is curving as much as possible within its speed limits set by spatiotemporal autocorrelation lengths, in order to control reaching movements.

We note that theorem (2) is not circular; i.e. it is not tautologically true that every possible measured neural state space dynamics, assuming enough neurons are recorded, will have dimensionality close to the NTC. In the supplementary material, we exhibit an analytical example of a very fast neural circuit, with a small temporal autocorrelation *τ*, recorded for a long time *T*, that nevertheless has dimensionality *D* much less that 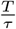 because the connectivity is designed to amplify activity in a small number of dimensions and attenuate activity in all others, similar to the way non-normal networks have been proposed to play a functional role in sequence memory [Ganguli et al., 2008b]. Finally, what kind of neural dynamics would have a maximal dimensionality, equal to its NTC? As we show in the Supplementary material, a random smooth manifold, with no other structure beyond smoothness, has such maximal dimensionality.

### 5 Beyond dimensionality: accurate recovery of the geometry of dynamical portraits

The above theory reveals a simple sufficient condition under which the dimensionality of dynamical portraits would remain unchanged if we recorded more neurons: namely if the number of recorded neurons exceeds the NTC, and the unrecorded neurons are statistically similar to the recorded neurons so as not to change population smoothness estimates. But importantly, even if we obtain the dimensionality of neural state space dynamics correctly by simply recording more neurons than the NTC, our theory so far does not provide any guarantee that we obtain their geometry correctly. Here we address the fundamental question of how many recorded neurons are sufficient to obtain the correct dynamical portrait of circuit computation at a given level of precision? By definition, the correct dynamical portrait is what we would obtain from dimensionality reduction applied to recordings of all the neurons in the behaviorally relevant brain region in question. Importantly, how does the sufficient number of recorded neurons scale with the complexity of the task, the desired precision, the total number of neurons in the brain region, and other properties of neural dynamics? And, interestingly, what minimal aspects of neural dynamics are important to know in order to compute this number?

To introduce our theory, it is useful to ask, when, intuitively, would recordings from a subset of neurons yield the same dynamical portrait as recordings from all the neurons in a circuit? The simplest visualizable example is a circuit of 3 neurons, where we can only measure 2 of them (Fig. 5A). Suppose the set of neural activity patterns encountered throughout the experiment consists of a single neural trajectory, that does not curve too much, and is somewhat randomly oriented relative to the single neuron axes (or equivalently, neural activity patterns at all times are distributed across neurons). Then the act of subsampling 2 neurons out of 3 is like looking at the shadow, or projection of this neural trajectory onto a coordinate subspace. Intuitively, it is clear that the geometry of the shadow will be similar to the geometry of the neural trajectory in the full circuit, no matter which 2 neurons are recorded. On the other hand, if the manifold is not randomly oriented with respect to single neuron axes, so that neural activity patterns may be sparse (Fig. 5B), then the shadow of the full neural trajectory onto the subspace of recorded neurons will not preserve its geometry across all subsets of recorded neurons. The challenge of course, is to make these intuitive arguments quantitatively precise enough to guide experimental design.

**Figure 5:**
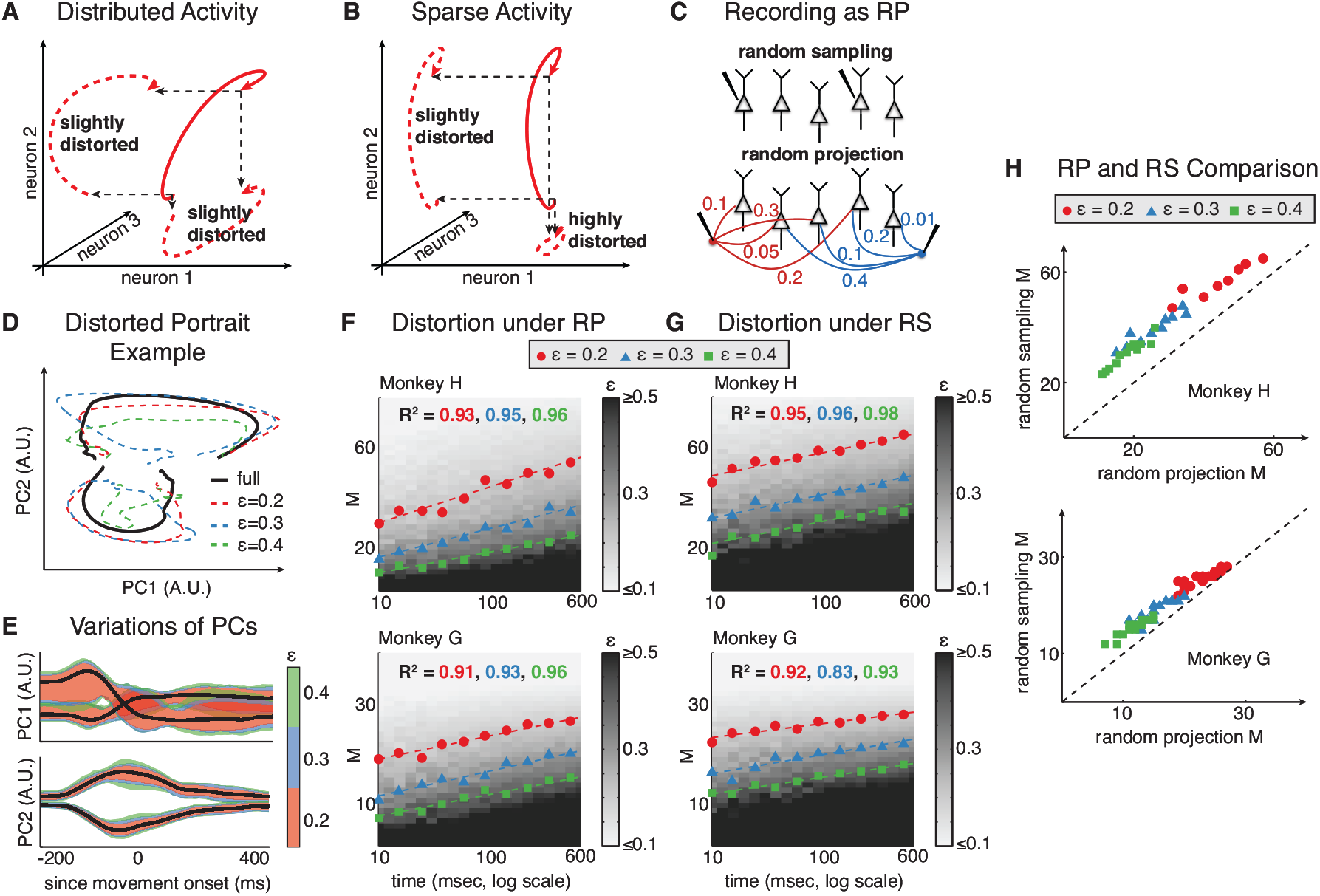
Conditions for accurate recovery of the geometry of dynamical portraits. **A**. A simple example of a *K* = 1 dimensional neural trajectory in *N* = 3 ambient dimensions, that is not aligned with respect to single neuron axes, so that neural patterns at all times are distributed across neurons. Recording any subset of *M* = 2 neurons induces only small geometric distortions in the neural trajectory relative to recording all *N* = 3 neurons. **B**. A simple example of a neural trajectory that is largely aligned with respect to the axis of neuron 2. Measuring any subset of neurons that does not include neuron 2 incurs a large distortion in the trajectory. **C**. For neural manifolds that are randomly oriented with respect to the single neuron axes, recording a random sample (RS) of *M* neurons (top) yields similar dynamical portraits as those obtained from recording a set of *M* random linear combinations, or random projection (RP), of all *N* neurons in the circuit. **D**. Example neural trajectories from two reach angles under different levels of distortion. The trajectories are projected into the top 2 PCSs, and only rotated for optimal alignment against fully observed trajectories. **E**. Variations (95% interval) of recovered PCs under different levels of distortion. **F**. For each monkey, the 95th percentile of the distortion distribution obtained after measuring *M* random linear combinations of all *N* recorded neurons (*N* = 109 for monkey H and *N* = 42 for monkey G), for random neural trajectory intervals of varying duration *T*. For each *M* and *T* we conducted 200 random trials and in 95% of trials the distortion *ϵ* between the resulting dynamical portrait and that obtained from all *N* recorded neurons for the same neural trajectory was less than the reported distortion. The iso-contrours of constant distortion are indeed straight lines in the *M* -log *T* plane, as predicted by Eq. (3). For the iso-contours *ϵ* = 0.2, 0.3, and 0.4, linear regression of *M* against log *T* yields excellent fits (*R*^2^’s of 0.93, 0.95 and 0.96 respectively for monkey H; 0.91, 0.93 and 0.96 for monkey G). **G**. Exactly the same analysis as **F** except now the measurement process corresponds to random samples of *M* neurons as opposed to *M* random projections of all *N* neurons. **H**. A quantitative comparison of panels *F* and *G* through a scatter plot of the number of randomly sampled neurons versus random projections required to obtain the same distortion on neural trajectories of varying duration.

To develop our theory of neural measurement, lets first assume the optimistic scenario in Fig. 5A, and pursue and test its consequences. If, in general, the neural data manifold (i.e. Fig. 3F) is randomly oriented w.r.t. the single neuron axes, then a measurement we can currently do, namely record from *M* randomly chosen neurons (Fig. 5C, top), becomes equivalent to a measurement we do not yet do, namely record from *M* random linear combinations of all neurons in the circuit (Fig. 5C, bottom). The former corresponds to projecting the neural manifold onto a coordinate subspace as in Fig. 5A, while the latter corresponds to projecting it onto a randomly oriented *M* dimensional subspace. If the neural manifold is randomly oriented to begin with, the nature of the geometric distortion incurred by the shadow, relative to the full manifold, is the same in either case.

This perspective allows us to then invoke a well-developed theory of how smooth manifolds in a high dimensional ambient space become geometrically distorted under a random projection (RP) down to a lower dimensional subspace [Baraniuk and Wakin, 2007, Clarkson, 2008]. The measure of geometric distortion *ϵ* is the worst case fractional error in euclidean distances between all pairs of points on the manifold, measured in the subspace, relative to the full ambient space (see Methods). The theory states that, to achieve a desired fixed level of distortion, *ϵ*, with high probability (*>* 0.95 in our analyses below) over the choice of random projection, the number of projections *M* should exceed a function of the distortion *ϵ*, the manifold’s intrinsic dimension *K* (1 for trajectories, 2 for surfaces, etc..), volume *V*, and curvature *C*, and the number of ambient dimensions *N*. In particular the theory states that 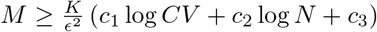, where *c*_1_, *c*_2_, and *c*_3_ are fixed constants, is sufficient. Thus intuitively, manifolds with low intrinsic dimensionality that do not curve much and have limited volume do not require that many measurements to preserve their geometry. Intriguingly, the number of ambient dimensions has a very weak effect; the number of required measurements grows only logarithmically with it. This is exceedingly fortunate, since in a neuroscience context, the number of ambient dimensions *N* is the total number of neurons in the relevant brain region, which could be very large. The logarithm thus ensures that this large number alone does not impose a requirement to make a prohibitively large number of measurements. Translating the rest of this formula to a neuroscience context, *K* is simply the number of task parameters, and *CV*, or curvature times volume, is qualitatively related to the NTC in (1); the numerator is the volume of the manifold in task space, and the reciprocal of correlation length is like curvature (short correlation length implies high curvature). Making this qualitative translation, the theory of neural measurement as a random projection suggests that as long as the number of recorded neurons obeys

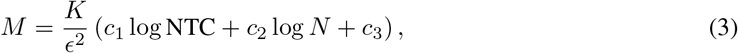

then we can obtain dynamical portraits with fractional error *ϵ*, with high probability over the choice of a random subset of recorded neurons. Remarkably, this predicts the number of recorded neurons need only scale logarithmically with the NTC to maintain a fixed precision.

Thus this theory makes a striking prediction that we can test in neural data: for a fixed number of task parameters *K*, and a fixed number of total neurons *N*, if we vary the number of recorded neurons *M* and the NTC, and compute the worst case fractional error *ϵ* in the recovered dynamical portraits relative to what we would get if we recorded all *N* neurons, then the iso-contours of constant distortion will be straight lines in a plane formed by *M* and the *logarithm* of the NTC. Of course we cannot currently record all *N* neurons in motor cortex, so we simply treat the dynamical portraits obtained in each monkey from all recorded neurons as the ground truth: i.e. we take *N* = 109 in monkey H and *N* = 42 in monkey G, as the total number of neurons, and we subsample a smaller number of neurons *M* from this pool. Also, we focus on the case *K* = 1, as the neural manifold in Fig. 3F is sampled smoothly only in time, and not angle; the reaches were done at only 8 discrete angles. Therefore we vary the NTC exactly as in Fig. 4C, by choosing random intervals of neural trajectories of varying durations *T* at each angle. For each interval duration *T*, which in this restricted context we can think of as simply proportional to the NTC, we use data from a random subset of *M* neurons, and compute the distortion *ϵ*(*M, T*) in the resulting dynamical portraits relative to the assumed ground-truth portrait obtained from all *N* recorded neurons. The theory above in Eq. (3) predicts exactly the same scaling in this scenario, with NTC replaced by time *T*.

Examples of the effects of different distortions *ϵ*, obtained by by sampling different sets of *M* neurons, are shown in Fig. 5D for dynamic portraits and Fig. 5E for individual PC’s. More generally, for each *M* and *T*, we conducted 200 trials and we plotted the 95’th percentile of resultant distribution of distortion *ϵ* as a heat map in Fig. 5FG. (i.e. 95% of trials had distortion less than what is reported). In panel F, we measured *M* random projections, or *M* random linear combinations of all *N* recorded neurons, for varying intervals of duration *T*, as the subsampled dataset, corresponding to the hypothetical experiment in Fig. 5C, bottom. It is clear that the iso-contours of constant distortion *ϵ* are well fit by straight lines in a plane formed by *M* and the logarithm of time *T*. This is a completely expected result, as this analysis is simply a numerical verification of an already proven theory. However, it forms a quantitative baseline for comparison in panel *G*, where we repeat the same analysis in panel *F*, except we record random subsamples of *M* neurons, as in experiment 5C, top. We obtain a qualitatively similar result as in panel *F*, which is remarkable, since this analysis is no longer a simple numerical verification of a mathematical theory. Rather, it is a stringent test of the very assumption that the neural manifold in Fig. 3F is randomly oriented enough with respect to single neuron axes so that random projections form a good theoretical model for the traditional measurement process of randomly sampling a subset of neurons. In essence it is a test of the assumption that the neural manifold is more like Fig. 5A than Fig. 5B, so that the two experiments in Fig. 5C yield similar geometric distortions in dynamical portraits as a function of recorded neurons *M* and neural task complexity NTC. In particular, the striking scaling of recorded neurons *M* with the logarithm of the NTC to maintain fixed precision in recovered dynamical portraits, predicted by the random projection theory of neural measurement, is verified. Moreover, we quantitatively compare the discrepancy between the two measurement scenarios in Fig. 5H, by creating a scatter plot of how many randomly sampled neurons versus random projections it takes to get the same distortion *ϵ* across all possible neural trajectory durations *T*, or equivalently, NTC’s. Even at a quantitative level, the data points are close to the unity line, relative to the total number of recorded neurons, suggesting that for this dataset, random projection theory is an impressively good model for the neural measurement process.

In the Supplementary Material, we study how these results are modified as neural activity patterns become more sparse and aligned with single neuron axes (i.e. less extreme versions of Fig. 5B). Remarkably, the linear scaling of number of neurons with the logarithm of NTC at fixed error is preserved, albeit with a higher slope and intercept. By comparing the neural data to simulated data with different levels of sparsity, we find that the neural data is indeed close to randomly aligned with respect to single neuron axes, as suggested by the closeness of the points in 5H to the unity line.

## 6 Discussion

### 6.1 An intuitive summary of our theory

Overall, we have generated a quantitative theory of trial averaged neural dimensionality, dynamics, and measurement that can impact both the interpretation of past experiments, and the design of future ones. Our theory provides both quantitative and conceptual insights into the underlying nature of two major order of magnitude discrepancies dominating almost all experiments in systems neuroscience: (1) the dimensionality of neural state space dynamics is often orders of magnitude smaller than the number of recorded neurons (e.g. Fig. 2), and (2) the number of recorded neurons is orders of magnitude smaller than the total number of relevant neurons in a circuit, yet we nevertheless claim to make scientific conclusions from such infinitesimally small numbers of recorded neurons. This latter discrepancy is indeed troubling, as it calls into question whether or not systems neuroscience has been a success or a failure, even within the relatively circumscribed goal of correctly recovering trial-averaged neural state space dynamics in such an undersampled measurement regime. To address this fundamental ambiguity, our theory identifies and weaves together diverse aspects of experimental design and neural dynamics, including the number of recorded neurons, the total number of neurons in a relevant circuit, the number of task parameters, the volume of the manifold of task parameters, and the smoothness of neural dynamics, into quantitative scaling laws determining bounds on the dimensionality and accuracy of neural state space dynamics recovered from large scale recordings.

In particular, we address both order of magnitude discrepancies by taking a geometric viewpoint in which trial-averaged neural data is fundamentally an embedding of a task manifold into neural firing rate space (Fig 3EF), yielding a neural state space dynamical portrait of circuit computation during the task. We explain the first order of magnitude discrepancy by carefully considering how the complexity of the task, as measured by the volume of the task manifold, and the smoothness of neural dynamics, as measured by a product of neural population correlation lengths across each task parameter, can conspire to constrain the maximal number of linear dimensions the neural state space dynamics can possibly explore. We define a mathematical measure, which we call neural task complexity (NTC), which, up to a constant, is simply the ratio of the volume of the task manifold and the product of neural population correlation lengths (Eq. (1)) and we prove (see Supplementary material) that this measure forms an upper bound on the dimensionality of neural state space dynamics (Eq. (2)). We further show in neural data from the motor cortex of reaching monkeys, that the NTC is much smaller than the number of recorded neurons, while the dimensionality is only slightly smaller than the NTC (Fig. 4). Thus the simplicity of the center out reach task and the smoothness of motor cortical activity, are by themselves sufficient to explain the low dimensionality of the dynamics relative to the number of recorded neurons. A natural hypothesis is that for a wide variety of tasks, neural dimensionality is much smaller than the number of recorded neurons because the task is simple and the neural population dynamics is smooth, leading to a small NTC. In such scenarios (experimental regime (ii) in Fig. 4A), only by moving to more complex tasks, not by recording more neurons, would we obtain richer higher dimensional trial averaged state space dynamics.

We address the second, more troubling, order of magnitude discrepancy by making a novel conceptual link between the time-honored electrophysiology tradition of recording infinitesimally small subsets of neurons in much larger circuits, and the theory of random projections, corresponding in this context to recording small numbers of random linear combinations of *all* neurons in the circuit. In scenarios where the neural state space dynamics is sufficiently randomly oriented with respect to single neuron axes (e.g. Fig. 5A) these two different measurement scenarios (Fig. 5C) yield similar predictions for the accuracy with which the dynamics are recovered as a function of the number of recorded neurons, the total number of neurons in the circuit, the volume of the task manifold, and the smoothness of the neural dynamics. A major consequence of the random projection theory of neural measurement is that the worst case fractional error in the geometry of recovered neural state space dynamics increases only with the *logarithm* of the total number of neurons in the circuit (Eq. (3)). This remarkable property of random projections goes hand in hand with why systems neuroscience need not be daunted by so many unrecorded neurons: we are protected from their potentially detrimental effect on the error of state space dynamics recovery by a strongly compressive logarithm. Moreover, the error grows linearly with the number of task parameters, and only again *logarithmically* with the NTC, while it decreases linearly in the number of recorded neurons (Eq. (3)). Thus recording a modest number of neurons can protect us against errors due to the complexity of the task, and lack of smoothness in neural dynamics.

This theory then resolves the ambiguity of whether systems neuroscience has achieved success or has failed in correctly recovering neural state space dynamics. Indeed it may well be the case that in a wide variety of experiments, we have indeed been successful, as we have been doing simple tasks with a small number of task parameters and NTC, and recorded in circuits with distributed patterns of activity, making random projections relevant as a model of the measurement process. Under these conditions, we have shown there is a modest requirement on the number of recorded neurons to achieve success. Our work thereby places many previous works on dimensionality reduction in neuroscience on much firmer theoretical foundations. Having summarized our theory, below we discuss its implications and its relations to other aspects of neuroscience.

### 6.2 Dimensionality is to neural task complexity as information is to entropy

To better understand the NTC, and its relation to dimensionality, it is useful to consider an analogy between our results and applications of information theory in neuroscience [Reike et al., 1996]. Indeed, mutual information has often been used to characterize the fidelity with which sensory circuits represent the external world. However, suppose that one reported that the mutual information rate 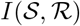 between the sensory signal 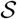 and the neural response 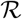 were 90 bits per second, as it is in the fly H1 neuron [Strong et al., 1998]. This number by itself would be difficult to interpret. However, just as dimensionality is upper bounded by the neural task complexity, mutual information *I* is upper bounded by entropy *H*, i.e. 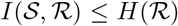. Thus if one measured the entropy rate of the response spike train to be 180 bits per second [Strong et al., 1998], then by comparing the mutual information to the entropy one could make a remarkable conclusion, namely that the neural code is highly efficient: the fidelity with which the response 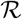 codes for the stimulus 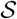 is within a factor of 2 of the fundamental limit set by the entropy of the neural response 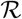.

Similarly, the observation that the dimensionality of recordings of 109 neurons in Monkey H in Fig. 4 is 7.1, is by itself, difficult to interpret. However, if one computed the NTC to be 10.5, then by comparing dimensionality to the NTC, one could make another remarkable conclusion, namely that motor cortex is exploring almost as many dimensions as possible given the limited extent, or volume of behavioral states allowed by the task, and the limited speeds with which neural population dynamics can co-vary across behavioral states. Thus just as entropy, as an upper bound on mutual information, allows us to measure the fidelity of the neural code on an absolute scale from 0 to 1 through the ratio of information to entropy, the NTC, as an upper bound on dimensionality, allows us to measure the complexity of neural state space dynamics on an absolute scale from 0 to 1, through the ratio of dimensionality to NTC. When this ratio is 1, neural dynamics is as complex, or high dimensional, as possible, given task and smoothness constraints.

### 6.3 Towards a predictive theory of experimental design

It may seem that neural task complexity could not be useful for guiding the design of future experiments, as its very computation requires knowing the smoothness of neural data, which would not have yet been collected. However, this smoothness can be easily estimated based on knowledge of previous experiments. As an illustration, consider how one might obtain an estimate of how many neurons one would need to record in order to accurately recover neural state space dynamics during a more complex reaching task in 3 dimensions. For concreteness, suppose a monkey has to reach to all points on a sphere of fixed radius centered at the shoulder of the reaching arm. The manifold of trial averaged task parameters is specified by time *t* into the reach, which varies from 0 to *T* ms, and the azimuthal and altitudinal angles *ϕ* and *θ*, each of which range from 0 to 2*π*. Now lets assume the smoothness of neural population dynamics across time will be close to the average of what we observed for 2 dimensional reaches (*τ* = 126 and 146 ms in monkeys H and G). Also lets assume reaches will take on average *T* = 600 ms as it did in the case of two dimensional reaches. Then the we obtain the estimate 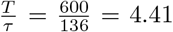. Now again, lets assume that both azimuthal (∆_*φ*_) and altitudinal (∆_*θ*_) neural correlation lengths would be the average of the angular correlation length of two dimensional reaches (∆ = 1.82 and 1.91 radians in monkeys H and G), yielding 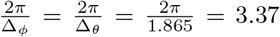. Then the NTC, according to Eq. (1), is proportional to the product of these 3 numbers, where in 3 dimensions, the constant of proportionality could be taken to be 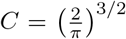. This product yields an estimate of 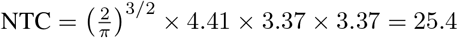.

If we trust this estimate, then this simple computation, coupled with our theorems proven above, allows us to make some predictions that can guide experimental design. For example, the theorem in Eq. (1) implies that no matter how many neurons we record in monkey motor and premotor cortices, the dimensionality of the trial averaged state space dynamics will not exceed 25. Moreover, the theorem in Eq. (3) tells us that if we wish to recover this state space dynamics to within fractional error *ϵ* = 0.2, relative to what we would obtain if we recorded all task relevant neurons, then we should record at least *M* = 300 neurons (see Supplementary Material). Now of course, we may not wish to trust this estimate, because we may have mis-estimated the neural correlation lengths. To be safer, we could easily underestimate the correlation lengths, and thereby obtain a safe overestimate of the NTC and requisite number of neurons to record. But overall, in this fashion, by combining an estimate of a likely NTC in future experiments with the new theorems in this work, we can obtain back of the envelope estimates of the dimensionality and accuracy of recovered state space dynamics, as neuroscience moves forward to unravel the neural basis for more complex behavioral and cognitive acts.

### 6.4 Departures from our assumption of statistical homogeneity

A critical assumption in using our theory to guide future experiments is that the set of unrecorded neurons is statistically similar to the set of recorded neurons, so that the denominator of the NTC in Eq. (1) will not change much as we fix the task and record more neurons. There are several important ways that this assumption could be violated. For example, there could be strong spatial topography in the neural code of the relevant circuit, so that as we expand our electrode array to record more neurons, the new neurons might have fundamentally different coding properties. Also, we may wish to record from multiple task relevant brain regions simultaneously, in which case our theory would have to apply to each brain region individually. Moreover, there may be multiple cell types in the relevant brain region. Unfortunately, most electrophysiology recordings do not give us access to cell type information (though spike width can sometimes serve as a proxy for the excitatory/inhibitory distinction). Thus the recovered neural state space dynamics reflects the combined action of all cell types. However, if we had access to cell type information, me may wish to define state space dynamical variables for each cell type. Then the theory would apply to each cell type alone. However, if cells of different types are strongly coupled, it is not clear that the collective dynamics of the circuit should be well explained by reduced degrees of freedom, or state, that are in one-to-one correspondence with cell types. This is an important empirical issue for further studies.

In essence, our theory applies to a spatially well mixed, statistically homogenous, localized brain region whose dynamics is relevant to the task. Fortunately, a wide variety of phylogenetically newer brain regions that evolved to learn new connectivities to solve tasks that evolution itself could not anticipate, for example prefrontal, parietal, pre-motor and motor cortices, and even older hippocampal circuits, exhibit precisely these kinds of mixed representations, in which the coding properties of individual neurons exhibit no discernible spatial topography, and almost every neuron codes for a mixture of multiple task parameters (e.g. [Machens et al., 2010, Mante et al., 2013, Rigotti et al., 2013, Raposo et al., 2014]). These are precisely the properties that make the neural manifold randomly oriented with respect to single neuron axes, and therefore make our random projection theory of neural measurement relevant, and the recovery of state space dynamics relatively easy despite subsampling.

But ironically, these very same properties make the goal of understanding what each and every individual neuron in the circuit does a seemingly difficult and questionable endeavor. While this endeavor has indeed traditionally been the putative gold standard of understanding, perhaps instilled in systems neuroscience by the tremendous success of Hubel and Wiesel in discovering single cell orientation tuning in primary visual cortex [Hubel and Wiesel, 1959], it is unclear that it will continue to be a profitable path going forward, especially in recently evolved brain regions where mixed representations dominate. But fortunately, the path of moving away from understanding single neurons to recovering collective state space dynamics, is a promising route forward, and indeed one that has firmer theoretical justification now, even in the face of extreme neural subsampling.

### 6.5 A why question: the neuroscientist and the single neuron

We have shown that when neural representations are distributed, or randomly enough oriented with respect to single neuron axes (Fig. 5A), so that random projections constitute a good model of the neural measurement process (Fig. 5C), then the life of the neuroscientist studying neural circuits becomes much easier: he or she can dramatically subsample neurons, yet still recover global neural state space dynamics with reasonable accuracy. However, neural systems evolved on earth long before neuroscientists arrived to study them. Thus no direct selection pressures could have possibly driven neural systems to self-organize in ways amenable to easy understanding by neuroscientists. So one could then ask a teleological question: why did neural systems organize themselves this way?

One possible answer lies in an analogy between the neuroscientist and the single neuron, whose goals may be inadvertently aligned. Just as a neuroscientist needs to read the state of a cortical neural circuit by sampling *O*(100) randomly chosen neurons, a downstream cortical neuron needs to compute a function of the state of the upstream circuit while listening to *O*(10, 000) neurons. Intuitively, if neural activity patterns are low dimensional enough, and distributed enough across neurons, then the single neuron will be able to do this. Indeed, a few works have studied constraints on neural representations in the face of limited network connectivity. For example, [Valiant, 2005] showed that the sparser neural connectivity is, the more distributed neural representations need to be, in order for neural systems to form arbitrary associations between concepts. Also [Sussillo and Abbott, 2012] showed that if neural representations in a circuit are low dimensional and randomly oriented with respect to single neuron axes, then a neuron that subsamples that circuit can compute any function of the circuit’s state that is computable by a neuron that can listen to all neurons in the circuit. And finally, [Kim et al., 2012] showed that the hippocampal system appears to perform a random projection, transforming a sparse CA1 representation of space into a dense subicular representation of space, yielding the ability to communicate the output of hippocampal computations to the rest of the brain using very few efferent axons.

These considerations point to an answer to our teleological question: in essence, our success as neuroscientists, in the accurate recovery of neural state space dynamics under extreme subsampling, may be an exceedingly fortunate corollary of evolutionary pressures for single neurons to communicate and compute accurately under the constraints of limited degree network connectivity.

### 6.6 Beyond the trial average: towards a theory of single trials

A natural question is how this theory would extend to the situation of single trial analyses. Several new phenomena can arise in this situation. First, in any single trial, there will be trial to trial variability, so that neural activity patterns may lie near, but not on, the trial averaged neural manifold, illustrated for example in Fig. 3F. The strength of this trial to trial variability can be characterized by a single neuron SNR, and it can impact the performance of various single trial analyses. Second, on each and every trial, there may be fluctuations in internal states of the brain, reflecting potentially cognitive variables like attention, or other cognitive phenomena, that are uncontrolled by the task. Such fluctuations would average out in the trial averaged manifold, but across individual trials would manifest as structured variability around the manifold. It would be essential to theoretically understand methods to extract these latent cognitive variables from the statistics of structured variability. Third, the trial averaged neural manifold may have such a large volume, especially in a complex task, so that in a finite number of trials *P* we may not be able to sufficiently cover this volume. One would like to know then, what is the minimum number of training trials *P* one would require, to successfully decode behavioral or cognitive variables on subsequent, held-out, single trials. Moreover, how would this minimum number of trials scale with properties of the trial averaged manifold, obtained only in the limit of very large numbers of trials?

We have already begun to undertake a study of these and other questions. Our preliminary results, some of which were stated in [Gao and Ganguli, 2015], suggest that the basic theory of trial averaged neural data analysis forms an essential springboard for addressing theories of single-trial data analysis. For example, in the case of the last question above, we find that the number of training trials *P* must scale with the NTC of the trial averaged neural manifold, in order for subsequent single trial decoding to be successful. Moreover, we have analyzed theoretically the recovery of internal states reflected in the spontaneous activity of large model neural circuits, while subsampling only *M* neurons for a finite amount of time 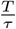 (a dimensionless ratio of recording time to single neuron correlation time *τ*). We find that the dynamics of these internal states can be accurately recovered as long as (a) both *M* and 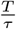 exceed the intrinsic dimensionality explored by the manifold of latent circuit states, and (b) the square-root of the product of neurons *M* and 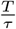 exceeds a threshold set by both this dimensionality and the single neuron SNR [Gao and Ganguli, 2015]. In turn the dimensionality of the manifold of latent circuit states is upper bounded by its NTC, so the NTC of a latent neural manifold determines the viability of single trial analyses, just as it does in the recovery of neural manifolds explicitly associated with externally measured task parameters. And finally, one may be tempted to conjecture that, due to finite SNR, single trial decoding performance may grow without bound as the number of recorded neurons increase - a result that is qualitatively different from the trial averaged theory, which suggests that only modest only numbers of neurons are required to accurately recover neural state space dynamics. However, there are several reasons to believe that such a qualitative discrepancy may not bear out. For example, neural noise may be embedded in the same direction as the signal, resulting in information limiting correlations [Moreno-Bote et al., 2014]. Moreover, empirically, in single trial decoding in the brain machine interface community, decoding performance already achieves a point of diminishing returns at even modest numbers of recorded neurons. The precise theoretical reasons for this remain an object of future study.

But overall, our initial results in a theory of single-trial analyses, to be presented elsewhere, suggest that the theory of trial-averaged neural dimensionality, dynamics and measurement, presented here, not only provides interpretive power for past experiments, and guides the design of future trial averaged experiments, but also provides a fundamental theoretical building block for expansion of the theory to single trial analyses. In essence, this work provides the beginnings of a theoretical framework for thinking about how and when statistical analyses applied to a subset of recorded neurons correctly reflect the dynamics of a much larger, unobserved neural circuit, an absolutely fundamental question in modern systems neuroscience. A proper, rigorous theoretical understanding of this question will be essential as neuroscience moves forward to elucidate the neural circuit basis for even more complex behavioral and cognitive acts, using even larger scale neural recordings.

## 7 Materials and Methods

### 7.1 Dimensionality Measure

Our measure of dimensionality is derived from the eigen-spectrum of the neuronal covariance matrix. This matrix underlies PCA, and indicates how pairs of neurons covary across time and task parameters (see Supplementary material). The eigenvalues of this matrix, *µ*_1_ ≥ *µ*_2_ ≥, …, *µ*_*M*_, reflect neural population variance in each eigen-direction in firing rate space. The participation ratio (PR),

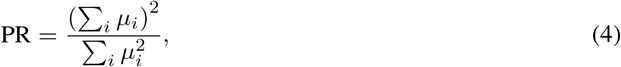

is a natural continuous measure of dimensionality. Intuitively, if all variance is concentrated in one dimension, so that *µ_α_* = 0 for *α* ≥ 2, then PR=1. Alternatively, if the variance is evenly spread across all *M* dimensions, so that *µ*_1_ = *µ*_2_ = … *µ*_*M*_, then PR= *M*. For other PCA eigenspectra, the PR sensibly interpolates between these two regimes, and for a wide variety of uneven spectra, the PR corresponds to the number of dimensions required to explain about 80% of the total population variance (see Supplementary material).

### 7.2 Preprocessing of the Motor Cortical Dataset

We use multi-electrode array recordings from two monkeys’ (H and G) PMd and M1 areas as they performed an eight-direction center-out delayed reach task [Yu et al., 2007]. There are between 145 and 148 trials in monkey H’s dataset and between 219 and 222 trials in monkey G’s dataset for each of the eight reach directions. Neural activities from each trial are time aligned to hand movement onset (time of 15% maximal hand velocity), and restricted to the −250ms to 350ms range time window around movement onset. Each spike train is smoothed with a 20ms gaussian kernel, and averaged with trials of the same reach angle to obtain the averaged population firing rates for the eight conditions. To homogenize the activity levels between neurons of different firing rates, and to highlight variability in the data resulting from task conditions, we further applied the square-root transform to population firing rates [Thacker and Bromiley, 2001], and subtracted their cross-condition average [Churchland et al., 2012].

### 7.3 Distortion Measure

To quantify the geometric distortion of dynamic portraits incurred from projecting them down from the *N* dimensional space of all neurons to the *M* dimensional subspace of recorded neurons (or *M* dimensional random subspace for a random projection), we adopt the pairwise distance distortion measure widely employed in the theory of random projections. Let **P** be the *M*-by-*N* linear projection operator that maps points from the full *N*-dimensional neural space into the *M*-dimensional subspace. For any pair of neural activity patterns **x**^*i*^ and **x**^*j*^ in the full *N*-dimensional space, the pairwise distance distortion induced by **P** is defined as

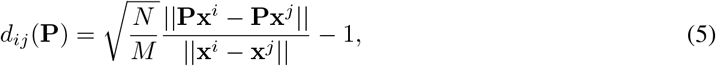

where the 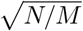 ratio compensates for the global distortion introduced simply by the reduction in dimensionality, and ║**v**║ denotes the Euclidean length of a vector **v**. A distortion of 0 indicates that the pairwise distance is the same both before and after the projection (up to an overall scale). The worst case distortion over all pairs of points (*i, j*) on the neural manifold is given by,

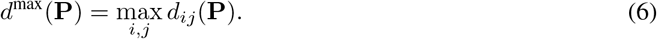

Since under either random projection or random sampling, **P** is a random mapping, *d*^max^(**P**) is a random variable. We characterize the distortion by the 95th percentile of the distribution of this random variable, i.e. that *ϵ* for which

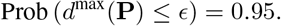

Thus with high probability (95%), over the random set of *M* measurements, the worst case distortion over all pairs of points on the neural manifold, will not exceed *ϵ*. In Fig. 5, for each value of *M* and *T*, we estimated *ϵ* by computing *d*^max^(**P**) 200 times for different random choices of **P**, and set *ϵ* to be the 95th percentile of this empirical distribution of distortions.

## 8 Acknowledgements

We thank S. Lahiri for inputs on dimensionality upper bound proofs. This work was supported by the Stanford Graduate Fellowship (P.G. and E.T.), the Mind, Brain and Computation Trainee Program (NSF IGERT 0801700, P.G. and E.T.), the Ruth L. Kirschstein National Research Service Award Predoctoral Fellowship (E.T.), NIH Director’s Pioneer Award (8DPIHD075623) and the Burroughs Wellcome, Simons, McKnight, and James S. McDonell foundations.

## Supplementary Materials: A Theory of Multi-Neuronal Dimensionality, Dynamics and Measurement

### I Introduction

In this supplementary material, we fill in the technical details associated with the main manuscript, and present proofs of the theorems stated there. The outline of the supplementary material is as follows.

In Sec. II, after reviewing principal components analysis, we justify the participation ratio as a reasonable measure of neural dimensionality, and explain its relation to more traditional measures like fraction of variance explained as a function of the number of principal components retained. We also explain rigorously how neural dimensionality is intimately related to task dimensionality through simple properties of linear algebra. Finally we compute the dimensionality of simple stationary tasks.

In Sec. III we introduce the logic behind neural task complexity (NTC) and why it places an upper bound on neural dimensionality. Along the way we prove a series of rigorous theorems in which we successively destroy structure in neural data, and show that the resultant destroyed dataset has higher dimensionality than the original dataset. The final outcome of this destruction of structure yields a simple dataset that is characterized purely by the number and range of task parameters and the local smoothness of the original dataset along each task parameter. The dimensionality of this simple dataset is by definition the NTC, and by construction it forms an upper bound on the dimensionality of the original dataset. Along the way we prove several intermediate results: (1) destroying long range correlations in a dataset increases its dimensionality; (2) making a dataset more homogeneous increases its dimensionality; (3) the best stationary approximation to a dataset has higher dimensionality than the original one; and (4) a particular factorization of a dataset across multiple task parameters has higher dimensionality than the original. Overall, this section proves the central results of Equations (1) and (2) in the main paper.

In Sec. IV we provide examples of the relationship between measured dimensionality, number of recorded neurons, and neural task complexity for a diverse set of theoretical models for datasets. We begin with a set of discrete stimuli, which we do not consider in the main paper, but include here for completeness. We then consider the case of random smooth neural manifolds, as an example of a model dataset in which actual dimensionality equals the neural task complexity. For both of these datasets, we show that the measured dimensionality initially grows with the number of recorded neurons, but then saturates to the value of the actual dimensionality as soon as the number of recorded neurons exceeds the actual dimensionality by a factor of about 10. This is consistent with the theory of random projections discussed later in which the preservation of geometric structure in data (including its dimensionality) to within a fractional error, requires a number of neurons that varies inversely with the error tolerance. Finally, at the end of Sec. IV we provide a theoretical neural network model that generates a dataset that lies deep within the elusive experimental regime (iii) in Figure 4A of the main paper. In this regime, the actual dimensionality is much less than the NTC (as well as the number of measured neurons), indicating that some neural network property is constraining dimensionality to a smaller value than that predicted by limited recording time and neural smoothness alone. In our example, this property is non-normal amplification which rapidly constraints network activity to a small number of neural activity patterns.

In Sec. V we begin with a self-contained review of seminal results in random projection theory. We then explore random projections of smooth manifolds in detail using simulations to quantitatively determine different constants of proportionality relating the number of required projections to the volume, smoothness, and ambient dimension of the manifold. These numerical simulations demonstrate that the constants of proportionality, which were not measured quantitatively in prior work, are not that large and are indeed *O*(1). Finally, we explore the effects of sparsity on neural activity, confirming that the essential theoretical prediction verified in Fig. 5 in the main paper, namely that the number of recorded neurons *M* need only scale logarithmically with the NTC in order to achieve a constant level of distortion, holds even when the data is quite sparse.

### II The relation between neuronal and task dimensionality

#### II.I Review of principal components analysis as a dimensionality reduction method

Trial averaged neural data is often described by an *M*-by-*N*_*T*_ data matrix, **X**. The *M* rows of **X** correspond to *M* recorded neurons, while the *N*_*T*_ columns of **X** correspond to all experimental task conditions - i.e. the columns could range over all combinations of time points and task parameter values occurring throughout the experiment. Each matrix element **X**_*ia*_ reflects the firing rate of neuron *i* in task condition *a*. For example, for a simple task unfolding over time as shown in Fig. 1 of the main paper, each column of **X** represents a pattern of firing rates across *M* recorded neurons at some specified time, and *N*_*T*_ is equal to the total number of sampled time points.

Principal components analysis (PCA), as a method of dimensionality reduction, starts from the *M* by *M* neuronal covariance matrix,

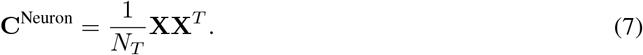

Here we have assumed that the data matrix **X** is centered, so that the sum of the columns is 0 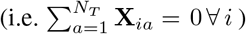. Geometrically, this means the cloud of *N_T_* neural activity patterns in *M* dimensional firing rate space, where each point in the cloud corresponds to a column of **X**, has its center of mass at the origin. Each matrix element 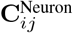 then reflects how strongly the firing rates of neurons *i* and *j* co-vary across task parameters. PCA relies on the eigen-decomposition of 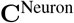, given by

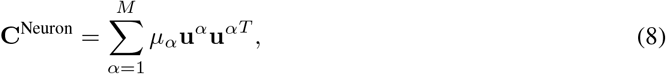

where **u**^*α*^’s are the eigenvectors of **C**^Neuron^ and the *µ*_*α*_’s denote the associated ordered eigenvalues, so that *µ*^1^ ≥ *µ*^2^ ≥, …, *µ*^*M*^. Each eigenvector **u**^*α*^ can be thought of as a static, spatial basis pattern of firing rates across neurons (i.e. the red and blue patterns in Fig. 1C of the main paper). Each eigenvalue *µ*_*α*_ reflects the amount of neural population variance along firing rate direction **u**^*α*^. Also, if we project neural activity onto pattern **u**^*α*^, we obtain the amount of pattern *α* in the neural population in task condition *a*:

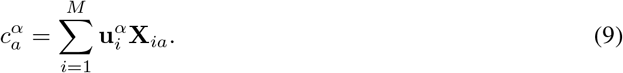

The variance of 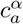 across task conditions *a* is precisely (up to a normalization by *N_T_*) the variance of the neural population in the direction **u**^*α*^, i.e. the eigenvalue *µ_α_*. In the special case, where *a* simply indexes time, we can think 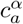 as one component of a dynamical trajectory in pattern space (i.e. the temporal components in Fig. 1C of the main paper). A low, *D* dimensional dynamic portrait of the trial-averaged data can then be obtained by plotting the top 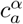 against each other, for *α* = 1*, …, D* (i.e. Fig. 1D of the main paper). The fraction of variance explained by the top *D* dimensions is 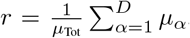, where 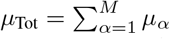 is the total variance in the neural population.

Dimensionality reduction by PCA is considered successful if a small number of patterns *D* relative to number of recorded neurons *M*, accounts for a large fraction of variance explained in the neural state space dynamics. One possible measure of dimensionality is the minimal number of basis patterns in the projection one must keep to achieve some pre-specified fraction, *r*, of the neural population’s total variance:

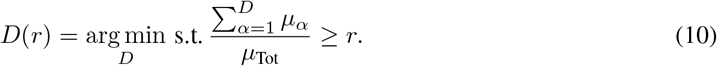

Note, that like any sensible measure of dimensionality based on the neuronal covariance eigenspectrum, this measure is invariant to an overall scaling of the eigenvalues. Fig. 2 of the main paper indicates that many experiments yield exceedingly low dimensional neural state space dynamics, in that a small number of dimensions relative to number of recorded neurons account for a large fraction of explained variance. Indeed, in many experiments, there is an order of magnitude discrepancy between dimensionality and number of recorded neurons, and one of our goals below is to understand the origin of such a large discrepancy, as well as to understand the accuracy of the dynamical portraits given by 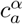 in the face of extreme subsampling in the number of recorded neurons.

#### II.II Participation ratio as a measure of dimensionality

While commonly used, measures of dimensionality associated with a given fraction of variance explained are not easily amenable to theoretical analysis. Here, we introduce a closely related measure, namely the participation ratio (PR) of the neural covariance eigenvalue spectrum, that obeys many of the same properties as more traditional measures, but is in contrast, much more amenable to theoretical analysis. The PR is defined as

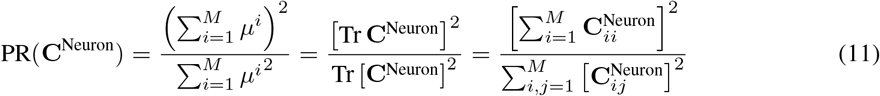

Like the fraction of variance explained measure, the participation ratio is also invariant to an overall scaling of eigenvalues. The PR is often used in statistical mechanics to quantify the number of active degrees of freedom in a thermally fluctuating system. Here we are using it to measure how many dimensions of the eigen-spectrum are active, relative to the total variance.

To gain an intuitive understanding of the participation ratio, let us consider an exactly rank-*D* neural covariance matrix **C**^Neuron^ with variance evenly spread across *D* dimensions: i.e. *µ*_*α*_ = *µ* for *α ≤ D*, and *µ*_*α*_ = 0 for *α > D*. The PR is invariant to the overall scale *µ* and evaluates exactly to *D*^2^*/D* = *D* as expected. For a more complex example, consider the eigenvalue spectrum of a typical trial-averaged dataset with full-rank but exponentially decaying eigenvalues: 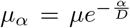. The spectrum’s PR is again independent of the overall scale *µ* and evaluates to 2*D* in the limit that *D ≪ M*, which is an intuitively sensible answer, and this value of dimensionality explains roughly 86% of the total variance. As we will further demonstrate below in more examples, the PR and the the measure of dimensionality *D*(*r*) based on a fraction *r* of variance explained, are often scaled versions of each other with the scaling constant depending on the shape of the eigenvalue spectrum. Moreover, for several typical spectra, keeping a number of dimensions equal to the PR leads to a fraction of variance explained between 80% to 90%.

While closely related to fraction of variance explained, the considerable theoretical advantage of the PR as a dimensionality measure is that it is an exceedingly simple function of the matrix elements of the neural covariance **C**^Neuron^. In particular, the numerator and denominator in Eq. (11) are simple quadratic functions of the matrix elements. In contrast the eigenvalues *µ*_*α*_ are highly complex functions of the matrix elements, as they are roots of the characteristic polynomial 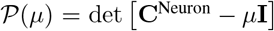, whose coefficients themselves are polynomials in the matrix elements of **C**^Neuron^ of all degrees ranging from 0 to *M*. In turn, *D*(*r*) defined in Eq. (10), as a nontrivial function of the eigenvalue spectrum, inherits this complexity as a function of the matrix elements of **C**^Neuron^. Thus the PR is a singular measure in that it embodies many of the intuitive properties we would associate with the notion of dimensionality, and is well correlated with traditional measures of dimensionality, yet it retains considerable analytical simplicity as a direct function of the matrix elements of **C**^Neuron^.

#### II.III A duality between neuronal dimension and task dimension

While the *M* by *M* neuronal covariance matrix **C**^Neuron^ is widely used for dimensionality reduction, and yields a set of neural basis patterns from which a dynamical portrait may be constructed, the overall structure or pattern of its matrix elements, 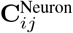 is usually nonintuitve and cannot be succinctly summarized. (See Supplementary Fig. 6A). Alternatively, consider the closely related *N*_*T*_ by *N*_*T*_ task covariance matrix,

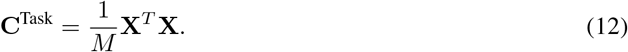

While the matrix elements 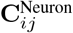 reflect how the firing rates of neurons *i* and *j* co-vary across *N*_*T*_ task parameters, the matrix elements 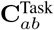 reflect how similar (if positive) or different (if negative) the neural population activity patterns across *M* neurons are for two different task conditions *a* and *b*.

**Supplementary Figure 6:**
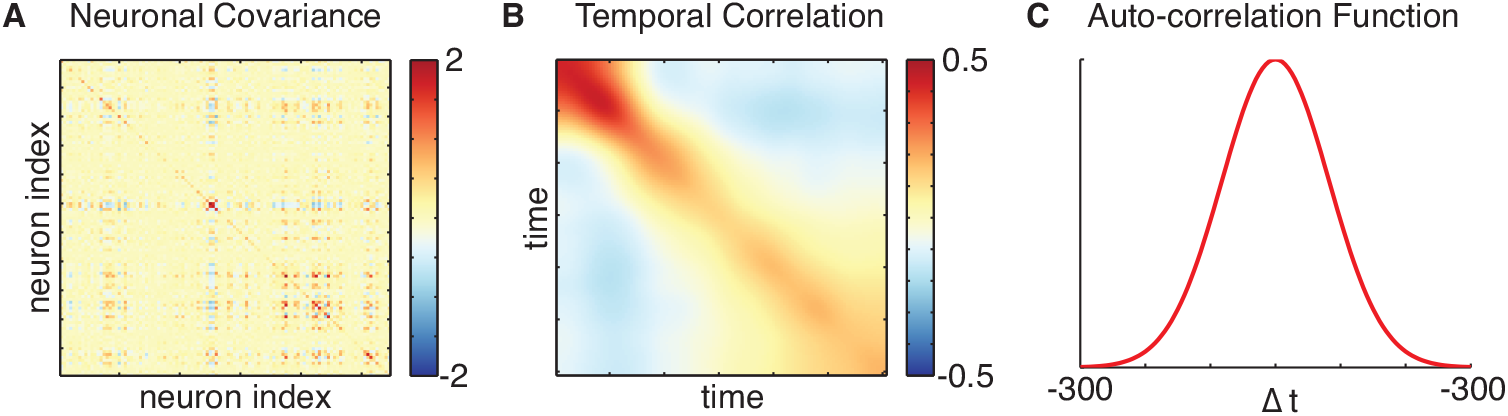
Neuron-by-neuron covariance versus task-by-task correlation. **A**. The complex *M* by *M* neuron-by-neuron covariance matrix for reaches to a single direction in Monkey H (*M* = 109 recorded neurons.) C^Neuron^, of the motor cortical data recorded from a monkey reaching to a single target; **B**. The corresponding *N_T_* by *N_T_* time-by-time task correlation matrix, C^Task^, where *N_T_* = 600, corresponding to the *T* = 600ms duration of a trial averaged reach with bin width 1ms. **C**. Example Gaussian auto-correlation function for a stationary task. Here the auto-correlation function is *f* (∆*t/τ*) where the time-scale *τ* = 125ms and *f*(*x*) is a Gaussian profile of variance 1.

A well known property of linear algebra [Strang and Aarikka, 1986] is that the nonzero eigenvalues of **C**^Task^ are exactly the same as the nonzero eigenvalues of **C**^Neuron^ (after a global rescaling of 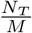 due to the specific normalizations chosen in Eq. (8) and (12)). This implies that any measure of dimensionality based on a scale invariant function of eigenvalue spectra (this of course includes both *D*(*r*) and PR), will be exactly the same, no matter whether it is computed with the spectrum of **C**^Neuron^ or **C**^Task^. It is this precise sense in which neural dimensionality and task dimensionality are inextricably linked.

The advantage of **C**^Task^ is that the overall structure or pattern of its matrix elements are often much easier to understand intuitively, and therefore to simplify. Compare for example Supplementary Fig. 6A to Supplementary Fig. 6B, as an example of a dual pair of **C**^Neuron^ and **C**^Task^ in the case of a simple task in which the only task parameter is time into a reach. While the former matrix elements are difficult to interpret, the latter are simple. They reflect the simple fact that patterns of neural activity are more similar to each other the more closely they occur in time relative to each other. When the neuronal dimensionality, measured through the PR, is computed using **C**^Task^ instead of **C**^Neuron^ in (11), it is clear that neural dimensionality is a very simple function of the similarity of neural activity patterns across pairs of task parameters. This is a key observation that will drive our theory below.

#### II.IV Dimensionality of simple stationary tasks unfolding over time

To gain further intuition behind the PR, and its relation to *D*(*r*), we consider the calculation of both in the special case of a simple task indexed only by time, where the task covariance matrix is *stationary* (up to boundary effects due to finite time *T*), which means that the similarity of two neural population activity patterns at two different times depends only on the relative time difference; i.e. 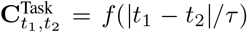, where *f*(*x*) is a smooth and symmetric auto-correlation function (Supplementary Fig. 6C). Here we have separated our description of the autocorrelation function into an intrinsic time scale *τ*, and an overall shape *f*(*x*) which takes as its argument a dimensionless ratio of actual time separation to *τ*. The width of *f*(*x*) as a function of *x* is chosen to be *O*(1).

Now, while in any given experiment, 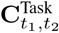 will never exactly be of this stationary form, we show below that we can always find a best stationary approximation to 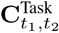, and that the dimensionality of this approximate stationary version of the task is higher than that of the original. Therefore, understanding the dimensionality of idealized stationary tasks becomes important, and also relevant to understanding the dimensionality of more general non-stationary tasks.

To begin, we first note that 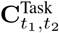 is of course a symmetric matrix, but also a Toeplitz matrix, as its diagonal elements 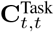 and off diagonal elements 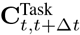, are independent of time *t*. As the size of such a *T* by *T* matrix becomes large, summations of any continuous function, *F*(·), of the matrix’s eigenvalues, like the numerator and denominator of the PR in Eq. (11), can be expressed using the Fourier transform of its central row [Gray, 1972], which is a discretized version of the auto-correlation function, *f*(*t/τ*). If we denote the Fourier transform of this autocorrelation by

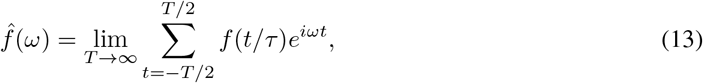

which exists as long as *f*(*x*) is absolutely summable, then we have

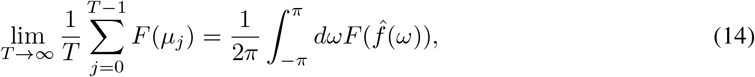

In practice, when the Toeplitz task-by-task correlation matrix is of a finite size *T*, the above expression still yields an accurate approximation under some restrictions: first, the stationary task needs to be in the regime where the duration of the task is much longer than the neural activities’ characteristic time scale, i.e. *T ≫ τ*. Second, when the recorded firing rates are discretized, binned or smoothed before analysis, the bin size or the smoothing kernel of the trial-averaged data needs to be much smaller than characteristic time scale. In other words, if *dt* is the temporal bin-width, then *τ ≫ dt* is needed for the approximation to be accurate.

Since both assumptions hold to a reasonable approximation in most large scale recordings collected in neuroscience today, we use Eq. (13) and Eq. (14) to compute the neural dimensionality of a stationary task. First, we reformulate the participation ratio in terms of the Fourier transform of the auto-correlation function governing the task-by-task correlation matrix,

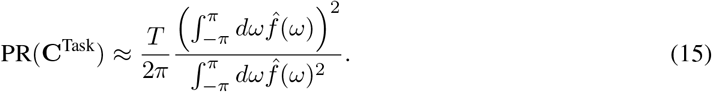

We then approximate the numerator term,

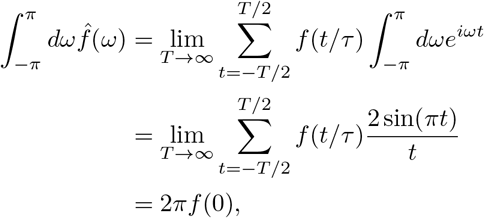

and the denominator term,

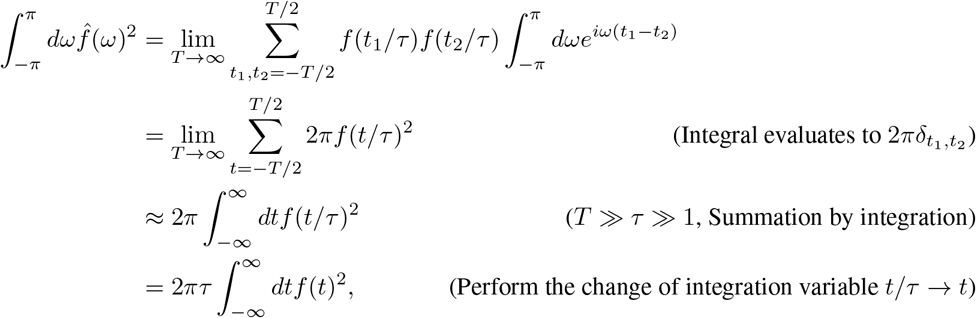

to obtain **C**^Task^’s participation ratio in terms of the task’s stationary auto-correlation function,

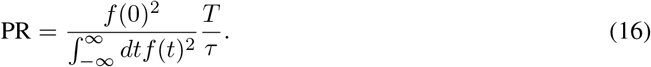

Interestingly, the measured dimensionality of neural state space dynamics associated with a simple stationary task scales linearly with the ratio of *T/τ*, with the constant of proportionality depending only on the shape of the auto-correlation function *f*(*x*). This result is the mathematical instantiation of the intuition embodied in Fig. 3C of the main paper, namely that a neural trajectory of duration *T* and autocorrelation with intrinsic timescale *τ* can explore at most order of magnitude 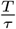 dimensions, because one must wait an amount of time *τ* before the neural trajectory can bend enough to explore another dimension. The constant of proportionality in Eq. (16) is, for a wide variety of autocorrelation profiles *f*(*x*), typical of neural data, simply an *O*(1) prefactor, which we explore next.

#### II.V Example stationary tasks: relation between participation ratio and fraction of variance explained

For concreteness, and further intuition, we compute the two measures of dimensionally, PR and *D*(*r*) for some example stationary tasks, with their respective auto-correlation functions. For the exponential, 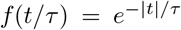, the Gaussian, 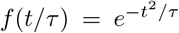 and the power-law, 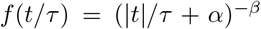, autocorrelation functions, we tabulate the analytical expressions for their participation ratio in Supplementary Table 1. In all three cases, their Fourier transforms are decreasing functions in frequency, which allows us to compute the dimensionality *D*(*r*) associated with a given fraction *r* of variance explained:

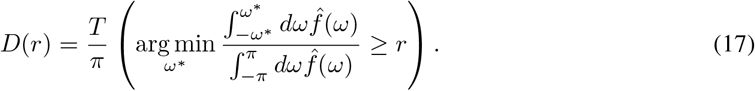

We obtain simple analytical expressions for *D*(*r*) in Supplementary Table 1 for the exponential and Gaussian auto-correlation functions. In these cases, just like the PR, *D*(*r*) also scales linearly with *T/τ* for any given desired *r*, indicating a semblance of universality in the conclusion that any reasonable measure of dimensionality is proportional to *T/τ* for a simple stationary task with intrinsic autocorrelation time scale *τ*. This holds true regardless of the particular fraction of variance *r* desired in the measure *D*(*r*), or the particular shape of the autocorrelations (exponential or Gaussian) we have examined.

**Supplementary Table S1:**
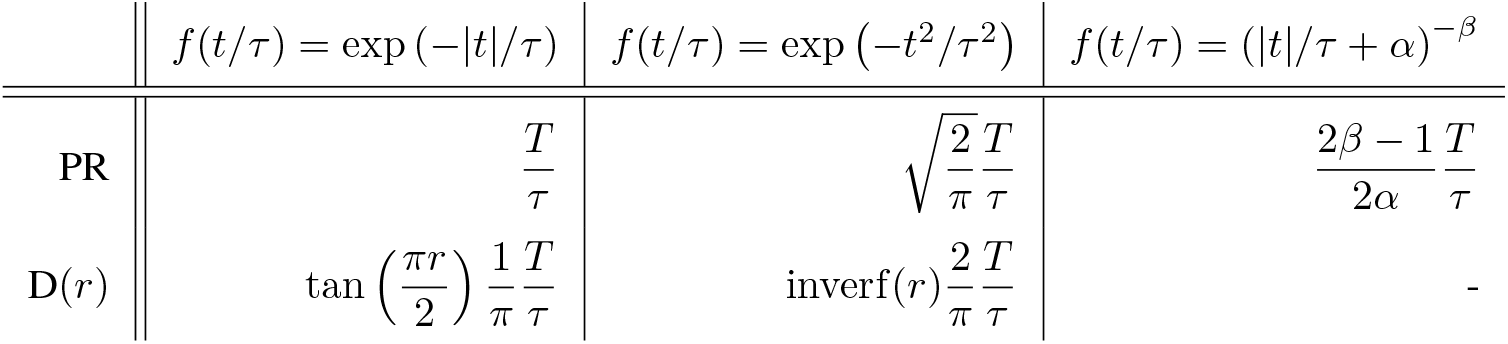
Analytical expressions for the participation ratio and the factional variance explained dimensionality measures for the Gaussian, exponential and power-law auto-correlation functions.

However, more quantitatively, what fraction of variance is actually explained by keeping a number of dimensions equal to the PR in these examples? This fraction *r* of variance explained corresponds to the solution to the equation *D*(*r*) = PR. For the exponential case, this equation reduces to 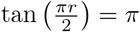, which has an approximate solution of *r* = 0.8. For the Gaussian case, this equation reduces to inverf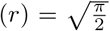, or 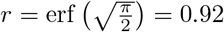.

We verify the correctness of these analytical expressions by comparing them against measured dimensionalities of numerically generated temporal correlation matrices under a variety of parameter combinations (Supplementary Fig. 7A,B). Overall, for both PR and *D*(*r*) (with *r* = 0.9), the numerical results show excellent agreement with our analytical expressions throughout the range of simulated *T/τ* values (Supplementary Fig. 2A,B). For the the dimensionality measure *D*(*r*) applied to the power-law auto-correlation function, the linear scaling with *T/τ* is numerically evident, despite the lack of an analytical formula.

**Supplementary Figure 7:**
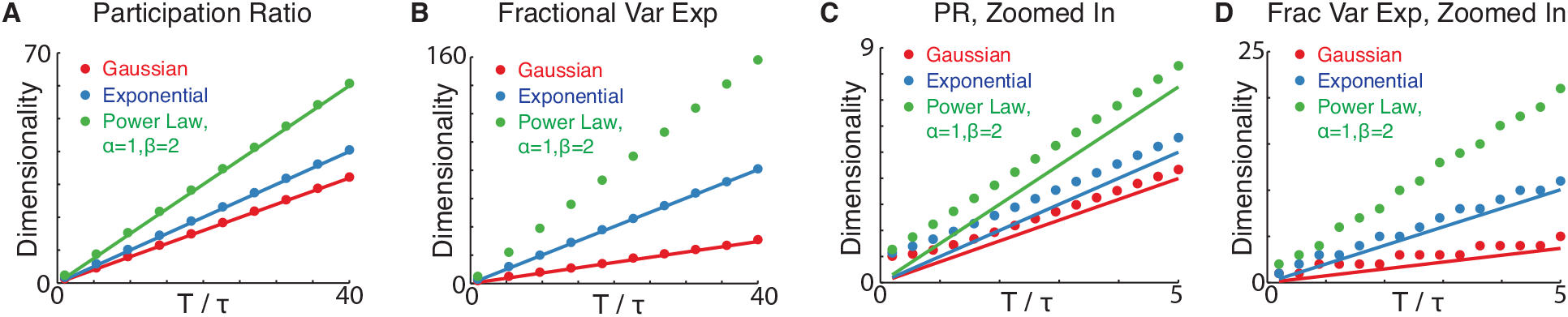
Numerical verifications of analytically computed participation ratios and fractional variance explained for the Gaussian, exponential and power-law auto-correlation functions. **A, B**. Analytical (lines) versus simulated (circles) participation ratio and fractional variance explained measured of dimensionality in the regime with large *T/τ*. **C, D**. Same as **A, B**, except zoomed in to show the small *T/τ* regime.

For completeness, we also conducted a comparison between the analytic approximations for dimensionality and their direct numerical calculation when *T/τ* is small, a regime in which the assumptions required for the accuracy of the analytical approximation to the eigenspectra of Toeplitz matrices in Eq. (14) are violated. As shown in the zoomed-in views (Supplementary Fig. 7C,D), the main deviation, in this regime, of the correct numerical result from its analytical approximation is a constant under-estimation by the analytical approximation. This deviation is simple to understand: if we consider a dataset with just two data points (so that the duration *T* equals the bin-width *dt*, which could be much less than the autocorrelation time *τ* - i.e. *T* = *dt ≪ τ*), the measured dimensionality will always be one, corresponding to the line connecting them, regardless of how close they are in time. The analytical expressions above for dimensionality are not valid in this regime, which violates the condition *dt ≪ τ ≪ T* necessary for accuracy of the analytic approximation to the eigenspectra of Toeplitz matrices in Eq. (14).

In summary, for a stationary task, characterized by an intrinsic autocorrelation time scale *τ* and total duration *T*, and sampled in time using bin width *dt*, both measures of dimensionality, *D*(*r*) for any *r* and PR, are proportional to *T/τ* in the experimentally relevant regime *dt ≪ τ ≪ T*, independent of the particular shapes of the auto-correlation function we have considered (exponential, Gaussian, and power law), and independent of the bin-width. Moreover, for a variety of typical auto-correlation functions, a number of dimensions equal to the PR explains about 80% to 90% of neural population variance in the data. Having now described the PR, and demonstrated its utility as a sensible measure of dimensionality, as well as its similarity to the more traditional measure *D*(*r*), we focus on the PR in our subsequent theoretical development, due to its theoretical simplicity.

### III Towards Neural Task Complexity: upper bounds on dimensionality through destruction of structure

We now aim to address the first order of magnitude discrepancy in systems neuroscience that we consider in this work, namely that the dimensionality of neural state space dynamics is often far less than the number of recorded neurons, as exemplified by the meta-analysis of 20 datasets in Fig. 2 of the main paper. We seek to explain the origin of this underlying simplicity. Our approach is to ask how high dimensional trial averaged neural state space dynamics could possibly be, given a limited extent of behavioral task parameters visited throughout any given experiment, and the smoothness of neural activity patterns across these task parameters. To address this, we describe a sequence of successive destructions of structure in neural state space dynamics, and prove that each destruction of structure necessarily increases the dimensionality of the dynamics. At the end point of this sequence of destruction, we are left with an exceedingly simple neural state space dynamics that has no structure whatsoever, above and beyond a limited volume and smoothness. The dimensionality of this resulting destroyed state space dynamics is by definition the neural task complexity (NTC) of the original state space dynamics, and by construction, this NTC constitutes an upper bound on the dimensionality of the original state space dynamics.

As described in the previous section, the structure of neural state space dynamics is well characterized by the *N*_*T*_ by *N*_*T*_ task correlation matrix **C**^Task^, and because of the duality between neural dimensionality and task dimensionality described above, we can compute neural dimensionality, as measured by its participation ratio, by replacing **C**^Neuron^ with **C**^Task^ in (11), obtaining,

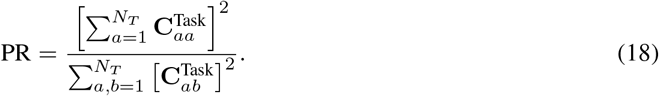

Any destruction of structure in the neural state space dynamics corresponds to a manipulation of the matrix elements 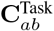, that essentially alters the correlation between the neural state, as a pattern of activity across *M* neurons, at two different task parameter values *a* and *b*.

#### III.I Increasing dimensionality by reducing long-range correlation

The simplest destruction of structure in **C**^Task^ is simply to reduce the magnitude of any off-diagonal element 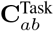 for *a* ≠ *b*. This decreases a single term in the denominator of (18) without changing the numerator, which only depends on the diagonal elements 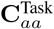, and therefore increases the dimensionality. Intuitively, reducing the squared correlation between neural activity patterns at two different task parameters allows the resulting dynamics to explore more dimensions.

As specific example, consider a single periodic neural trajectory that is confined to oscillate with period *τ*. Then the task parameter *a* indexes time *t*, and the task correlation matrix 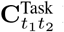 has peaks in off-diagonal bands whenever *t*_1_ − *t*_2_ is an integer multiple of *τ*, reflecting the fact that after every time *τ*, the neural dynamics is forced to revisit the same neural activity pattern by virtue of the oscillation. Consider instead a destroyed dynamics in which all off-diagonal elements 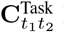 areset to 0 for all 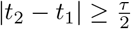. The resulting destroyed dynamics only has a central band of correlation for 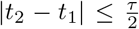, reflecting the smoothness in time, but not the periodicity, of the original dynamics. The destroyed dynamics behaves more like a smooth random walk with an approximate autocorrelation time *τ*, that never revisits the same point in neural state space over long times, and therefore can explore more dimensions than the original periodic dynamics. In this sense, the destruction of long-range temporal correlations necessarily increases dimensionality.

#### III.II Increasing dimensionality by homogenizing dynamics

A second way to destroy structure in the task-by-task correlation matrix 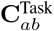 is to homogenize it by replacing each of a subset of the matrix’s off-diagonal entries with their average. This replacement increases dimensionality by again reducing PR’s denominator, without affecting the numerator, which only depends on diagonal elements. To see this, first consider Jensen’s inequality, which states that for any convex function *g*,

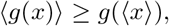

where *x* is any random variable drawn from a distribution *P*(*x*), and 〈·〉 denotes an average with respect to this distribution. To apply Jensen’s inequality, consider a specific subset of *K* off diagonal elements of 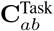, indexed by *γ* = 1, …, *K*, taking the values *x*_*γ*_. Here each *γ* indexes a particular pair of task parameters (*ab*) with *a ≠ b*, and *x*_*γ*_ is the value 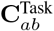. Let the distribution 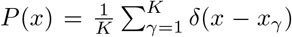. *P*(*x*) places equal probability mass on the chosen elements. Also let *g* be the convex function *g*(*x*) = *x*^2^. Then according to Jensen’s inequality, the average of the squares of *x*_1_, …, *x_K_* is greater than or equal to their average squared, or

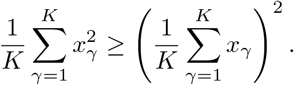

If we denote the average by 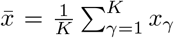, this means that replacing each *x*_*γ*_ with the average value 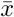 leads to a smaller denominator in Eq. (18), since the above inequality implies 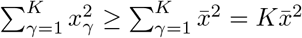. By reducing the denominator of the PR without changing the numerator, the dimensionality goes up.

Thus intuitively, any inhomogeneities in the off diagonal task-task correlation matrix 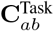 reflect inherent structure in the neural state space dynamics, through different degrees of similarity between neural activity patterns at different pairs of task parameters *a* and *b*. Destroying such structure by homogenizing it, i.e. replacing a subset of off-diagonal elements by their average, necessarily increases the dimensionality of the resulting destroyed dynamics.

#### III.III The best stationary approximation to a task has higher dimensionality than the original task

A simple, but important application of the above method for destroying structure is the result that the best stationary approximation of the task-by-task correlation matrix associated with a single task parameter (Supplementary Fig. 8A,B) has increased dimensionality. Consider a simple task where the task parameter *a* indexes time *t* only. In general such a task will be non-stationary, so that the matrix elements 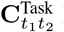 will not be a function of the time separation |*t*_2_ − *t*_1_| only. What is the best stationary approximation to this neural dynamics, described by a corresponding stationary task-task covariance matrix 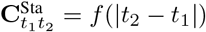, where by definition, the best one minimizes the squared error

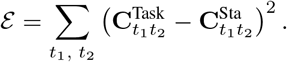

By the usual result that the single number 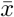 that minimizes the squared distance to a fixed collection of *K* numbers *x*_1_, …, *x*_*K*_ is the average of those numbers, 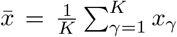, it is easy to see that the best stationary approximation is the one that averages the diagonals and off-diagonals of C^Task^:

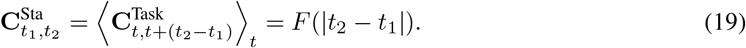

When *t*_2_ ≠ *t*_1_ the average corresponds to homogenizing in time an off-diagonal band of **C**^Task^, which by the above section increases dimensionality by decreasing the denominator of the PR in (18). However, the diagonal band is also averaged. Fortunately, this does not change the numerator of the PR, which is simply the sum of the diagonals squared. Thus overall, the best stationary approximation to a task, by destroying structure in the original task associated with temporal inhomogeneities, has increased dimensionality relative to the original task. See Supp. Fig. 8AB for an example of a non-stationary task and its best stationary approximation. In neural data analyzed in this work, this best stationary approximation is very well modeled by a Gaussian shaped autocorrelation profile (Supp. Fig. 8C).

**Supplementary Figure 8:**
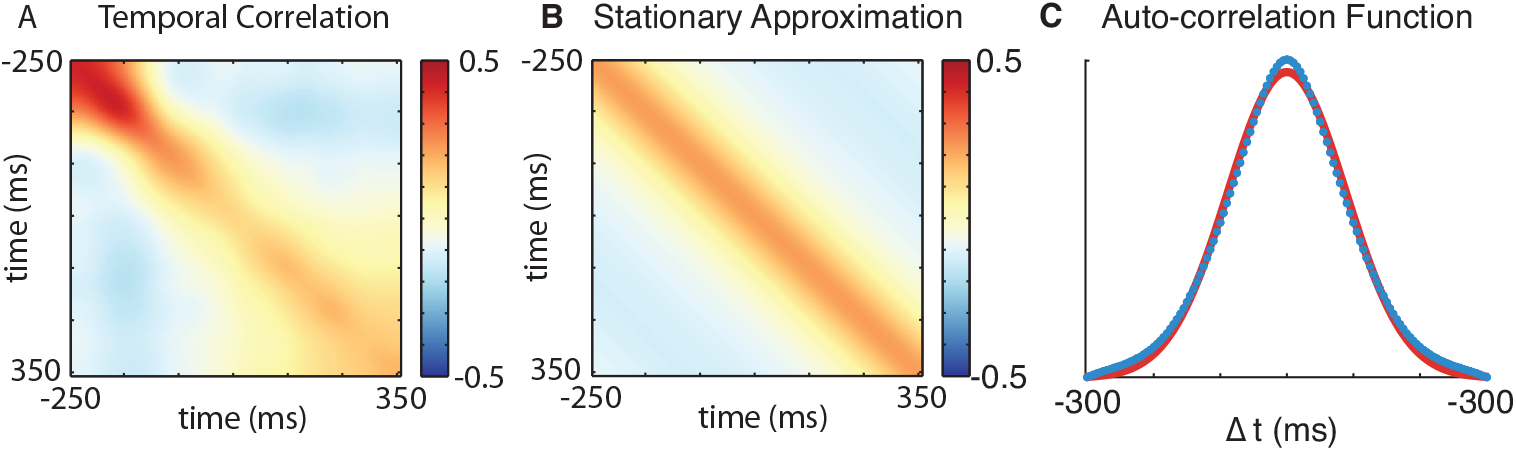
Example stationary approximation of a trial-averaged dataset. **A**. The task-by-task correlation matrix, C^Task^, of motor cortical state space dynamics while Monkey H is reaching to a single target. **B**. The best stationary approximation C^Sta^ to C^Task^ obtained by replacing the diagonal and the off-diagonals with their respective averages. **C**. A Gaussian fit (red) to the measured auto-correlation function of the best stationary approximation (blue).

In general, once we have the best stationary approximation **C**^Sta^ and its associated autocorrelation function *F* (∆*t*), we can proceed in one of several ways to obtain an upper bound on the dimensionally of the original neural state space dynamics described by **C**^Task^. First, a natural further destruction of structure is to destroy long range correlations, as described above, by setting to 0 *F*(∆*t*) for |∆*t*| greater than some threshold, which is chosen to include the central peak of *F* near the origin, but exclude other structure associated with peaks in *F* away from the origin. This process is designed to obtain dimensionality upper bounds that incorporate local smoothness alone, and no other longer range structure.

With this modified *F* one could write *F* in the form *F*(∆*t*) = *f*(∆*t/τ*), thereby separating the width of the central peak of *F*, as described by *τ*, and the shape of the central peak of *F*, as described by the function *f*(*x*) which has width in its dimensionless argument *x* of *O*(1). Then the resulting dimensionality upper bound is given by Eq. (16). This dimensionality upper bound is proportional to *T/τ*, up to a constant *O*(1) factor that depends on the detailed shape of *f*(*x*). It turns out that in the neural data analyzed in this paper, the shape of *f*(*x*) was well approximated by a Gaussian profile, yielding the constant factor 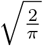 and the resulting NTC for a single task parameter of

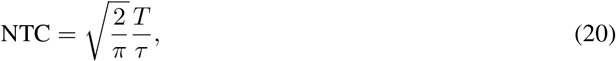

reported in the main paper.

Now this procedure for obtaining an NTC that upper-bounds neural dimensionality seems to depend on knowing many details of the neural data beforehand. However, it is exceedingly easy to estimate a likely NTC *before* the neural data is collected. Basically, one simply needs to estimate *τ*, the approximate width of the central peak of the temporal autocorrelation function of the neural trajectory, and know the total duration *T* of the neural trajectory. One need not know in detail the exact shape of the central peak, as this only contributes an *O*(1) constant of proportionality. For example, if it is Gaussian then this constant is 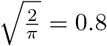, while if it is exponential then it is 1 (see e.g. Supp. Table 1). However, when one is attempting to explain 1 to 2 order of magnitude discrepancies between neural dimensionality and number of recorded neurons, then these *O*(1) differences in NTC due to different shapes of the auto-correlation function are not that important. Similarly, when one is attempting to estimate a likely NTC in future recordings for the purposes of designing experiments, these *O*(1) differences again are not as important - i.e. they may not be the dominant source of estimation error.

#### III.IV For multiple task parameters, a factorized stationary approximation to a task has higher dimensionality

For datasets with *K* task parameters, the task-by-task correlation matrix has a block structure. For example, for the eight-direction center-out reach task in shown in Figure 3D of the main paper with *K* = 2, the correlation matrix consists of eight-by-eight blocks of *T*-by-*T* temporal correlation or cross-correlation matrices (Supplementary Fig. 9A): entries of the (1, 1) block denote the correlations between neural activity patterns at different time points for reaches to the first target, whereas entries of the (1, 2) block denote the cross-correlations between neural activity patterns at some time during the reach to the first target and at a different time during the reach to the second target.

To upper bound the dimensionality of such compound correlation matrices, with multiple task parameters, we destroy structure in a specific way that leads to a stationary and factored approximation to the original correlation matrix. The dimensionality of this stationary factored approximation is then defined to be the NTC, and by construction, it upper bounds the dimensionality of the original data. To arrive at the stationary factorized approximation, we follow a two-step procedure (Supplementary Fig. 9B). The first step is to obtain a stationary approximation to the original task covariance matrix, using an extension of the logic we used in the *K* = 1 parameter case. With *K* parameters, the task manifold in parameterized by a *K*-tuplet of numbers (*t*^1^, … *t*^*K*^). Let two *K*-tuplets, 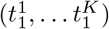 and 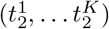 denote two points on the task manifold, and let the task correlation matrix 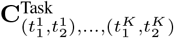 denote the correlation between the two neural activity patterns associated with two points on this task manifold. In general, this task correlation will not be stationary, in that it will not be a function of only differences between points, i.e. of 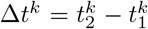.

We can find the best stationary approximation by averaging along off-diagonals through

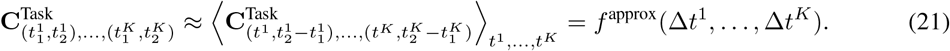

Via arguments similar to those presented in the previous section, the resulting stationary, block Toeplitz approximation of the task correlation matrix has a higher dimensionality than the original task correlation matrix. Interestingly, this approximation is diagonalizable, since it is an example of a Hewitt block Toeplitz matrix, but its correlation structure is still too complex for the resulting dimensionality upper bound to be intuitive and interpretable.

**Supplementary Figure 9:**
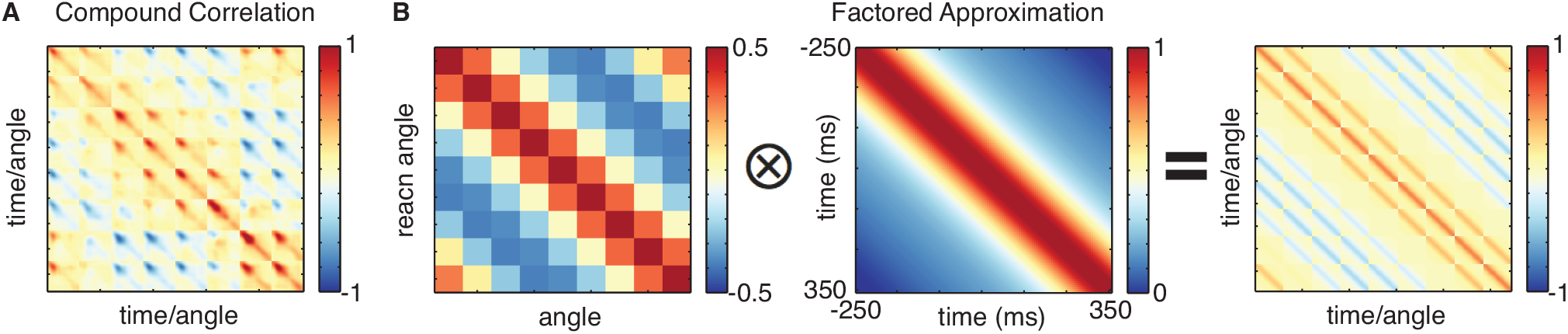
Example stationary and factor approximation of a trial-averaged dataset of a monkey doing the eight-direction center-out reach task. **A**. The compound task-by-task correlation matrix, C^Task^, of the motor cortical data during reaches to all possible directions. Inner indexes denote time during the reaches, while the outer blocks denote the different reach targets. **B**. The factored and stationary approximation to **A** as the Kronecker product of an angle-by-angle and a time-by-time stationary correlation matrices.

Instead, we apply a second approximation step to factorize the contributions of each task parameter. Formally, we decompose the approximate stationary compound correlation function, *f*^approx^(∆*t*^1^, …, ∆*t*^*K*^), into a product of individual task parameters’ auto-correlation functions,

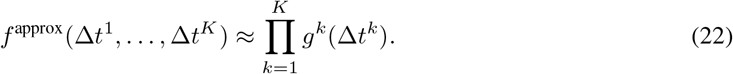

This factorization is the same as approximating the stationary compound correlation matrix with a Kronecker product,

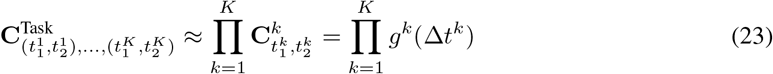

where each of the component matrix is a task parameter’s stationary correlation matrix, whose NTC and dimensionality is given by the corresponding auto-correlation function, *g*^*k*^(*·*). Since the resulting eigenvalues of the Kronecker product are all possible outer products of the form of 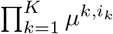, with *µ*^*k,i*_*k*_^ denotes the *i*_*k*_th eigenvalue of the **C**^*k*^ factor, the participation ratio of the factored and stationary approximation is,

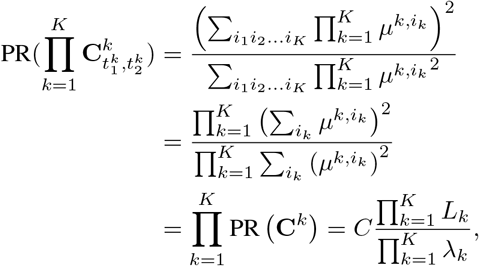

which is precisely the product of the NTCs for each of the task parameters.

For this approximation to be an upper bound on the original correlation matrix’s dimensionality, we still need to prescribe a factorization procedure that yields a guaranteed increase in dimensionality. The factorization we choose is a recursive projection procedure, where we successively replace the off-diagonal blocks of the stationary approximation with their projections onto the diagonal blocks. In terms of the auto-correlation factors, *g^k^*(·)s, the recursive projection procedure is given by,

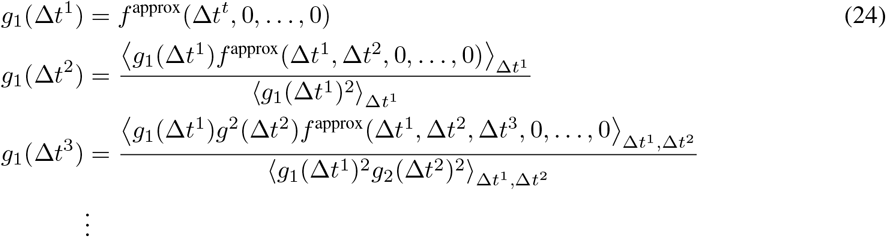

To see that this procedure produces an approximation that upper bounds dimensionlality, we first note that the numerator of the resulting approximation’s participation ratio is unchanged from the stationary approximation, since only the off-diagonal blocks are replaced by projections. Secondly, we note that the denominator of the participation ratio, which is the sum of squares of the off-diagonal blocks’ entries, can only be decreased by the projections. Consequently, the dimensionality of the factored approximation upper bounds that of the stationary approximation which in turn upper bounds that of the original task correlation matrix. For example, one can compare the original task correlation matrix (Fig. 9A) to its stationary factorized approximation (Fig 9B) in the case of a center out reach task studied in the main paper.

#### III.V Algorithmic computation of neural task complexity

Here we summarize the algorithmic steps required to compute the NTC.

To compute the NTC of a trial-averaged dataset with a single task parameter, *t*, in practice, we have the following procedure:

- Mean-subtract each neurons’ averaged activity across task parameters from each row of the neurons by task parameter data matrix **X**.
- Compute data’s task-by-task correlation matrix **C**^Task^ = **X**^*T*^**X**.
- Find the best stationary approximation of **C**^Task^, and construct the corresponding auto-correlation function 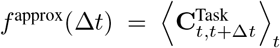. If necessary, set any long-range correlations to zero, while keeping the short-range correlations induced by smoothness alone.
- If *f*^approx^(·) can be fitted with a simple form, such as the Gaussian, exponential or power-law autocorrelation function whose participation ratio we’ve computed analytically (See Supplementary Table 1), extract the relevant auto-correlation scale *τ* and substitute it into the appropriate participation ratio expression to obtain the NTC.
- If the stationary approximation has a non-classical functional form for the auto-correlation function, one can still compute an NTC that depends, up to an *O*(1) factor, on the detailed shape of this auto-correlation function through the formula for the PR derived in (16).

To compute the NTC of a dataset with multiple task parameters, we have the following procedure:

- Mean-subtract each neurons’ activity averaged over all task parameters for rows of the data matrix **X**.
- Compute the compound task-by-task correlation matrix, **C**^Task^ = **X**^*T*^**X**.
- Compute **C**^Task^’s stationary approximation with the averaging procedure,

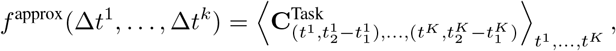

and its factorization with the prescribed recursive projection procedure (24),

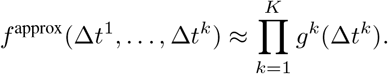
- Compute the correlation lengths and the dimensionalities for the individual task parameter’s the stationary approximation, **C**^*k*^, as described for the single-parameter case. The resulting product of dimensionalities is then the NTC for this dataset of multiple task parameters.

### IV Examples of the relationship between dimensionality and neural task complexity

#### IV.I Dimensionality and neural task complexity for discrete stimuli or behavioral conditions

Thus far, we focused on continuous task parameters that evoke smoothly evolving neural activity patterns, and showed how to compute a simple NTC measure that upper bounds the dimensionality of the neural data by taking into account only the extent of each task parameter and the neural correlation length along each task parameter. Is there a similar concept of NTC for stimuli or behaviors that are discrete? With *P* discrete stimuli, the resulting trial-averaged neural activity patterns can, of course, span a *P*-dimensional subspace of the neural space as long as the number of recorded of neurons exceeds that of the number of trials, i.e. *M ≥ P*. Thus *P* is a natural, if trivial, measure of NTC in this case.

A more interesting quantity in this situation may be the dimensionality of a *random* data set. Consider for example a situation in which *P* stimuli or behavioral conditions, indexed by *a* = 1, …, *P*, yield *P* random (mean subtracted) activity patterns across all *N* neurons in a circuit. By this we mean that the mean subtracted activity **x**_*ia*_ of neuron *i* in stimulus *a* are uncorrelated across both neurons and stimuli, and drawn i.i.d from a Gaussian distribution of mean 0 and variance 1 (since our dimensionality measures are independent of overall scale, we simply normalize the variance to be 1). This means that for large *N ≫ P*, the task correlation matrix of the full neural population, 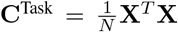, where **X** is the *N* by *P* data matrix, converges to a *P* by *P* identity matrix, and therefore has a dimensionality, measured by its PR, to be *P*. This is a reflection of the fact that in very high dimensions (*N ≫ P*), all 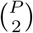 pairs of random vectors in *N* dimensional space are overwhelmingly likely to be close to right angles with respect to each other. Thus the *P* neural patterns form an orthogonal basis and have dimensionality *P*.

However, the situation is very different if we don’t record all *N* neurons, but instead record only *M* neurons, where *M* may be the same order of magnitude as the number of stimuli/behaviors, *P*. The data matrix **X** is then a smaller *M*-by-*P* dimensional random matrix, whose entries are generated i.i.d. with zero mean and unit variance. In the limit in which *M* and *P* are large, while the ratio is *O*(1), the distribution of eigenvalues of **C**^Task^, converges to the Marchenko-Pastur law [Marchenko and Pastur, 1967],

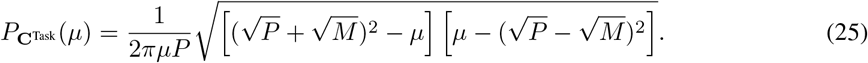

This law allows us to evaluate the numerator of the PR, which is the squared sum of the eigenvalues of **C**^Task^, as well as the denominator, which is the sum of squares of eigenvalues of **C**^Task^. The resulting dimensionality, or ratio of these two quantities, is simply,

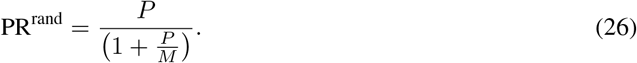

This of course reduces to the previous result of dimensionality *P* when the number of recorded neurons *M* is much larger than *P*. For example, if *M* = 10 × *P*, then the measured dimensionality would be about 0.9 × *P*. However, the cloud of neural activity patterns becomes geometrically distorted as we subsample neurons down to the point where *M* is not much larger than *P*. In that case, all 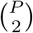 pairs of neural activity patterns are no longer sufficiently orthogonal, leading to a proliferation of small, but non-negligible, off diagonal elements in the *P* by *P* matrix **C**^Task^ that conspire to reduce the measured dimensionality of the random neural dataset, relative to what one would find if one recorded all *N* neurons.

The dimensionality PR^rand^ of a random dataset yields an interesting null model with which to compare dimensionality measured in actual experiments with *P* discrete stimuli, especially after the neural activity patterns have been normalized to have the same length in firing rate space. Indeed, comparison of measured dimensionality to this null model dimensionality, as opposed to the NTC upper bound, which is simply *P*, provides a powerful guideline for the interpretation of neural data. When the measured dimensionality is far below PR^rand^, the dataset is more correlated and lower-dimensional than what one would expect from random data, suggesting that circuit dynamics may be constraining neural activity patterns to live in a low dimensional space. On the other hand, when the measured dimensionality is higher than PR^rand^, the neural circuit may be actively de-correlating neuronal activity patterns to make them more orthogonal than one would expect by chance.

#### IV.II Random smooth manifolds have dimensionality equal to neural task complexity

We return to the case of a smooth, trial averaged neural state space dynamics in a task with *K* task parameters. What kind of state space dynamics would have a dimensionality that would necessarily saturate the upperbound set by the NTC, given we have recorded enough neurons *M*? Intuitively, the answer is a smooth random manifold that has the same autocorrelation lengths across task parameters and moreover factorizes across task parameters, with no other structure to further constrain its dimensionality. Such a random neural manifold for the state space dynamics is generated randomly by drawing each neuron’s response given the task parameters independently from a stationary random gaussian process with covariances determined by the data’s factored correlations. Formally, the joint distribution of the neural activities is Gaussian, and obeys the following,

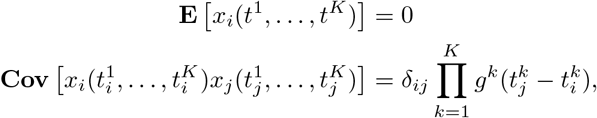

where *x_i_*(*t*^1^, …, *t*^*K*^) denotes neuron *i*’s firing rate at the point in the task manifold given by the *K*-tuplet (*t*^1^, …, *t*^*K*^). Since the task-by-task correlation matrix of this generated manifold is exactly factored and stationary as specified, its measured dimensionality must saturate the NTC upper bound constrained by only smoothness, as long as we measure enough neurons *M*, so that the empirically measured autocorrelation functions converge in limit to those that generated them.

We simulate such a smooth random manifold with *K* = 2 and gaussian auto-correlation functions with circular boundaries parameterized by lengths *L*^1,2^ = 100 and autocorrelation lengths *λ*^1,2^ = 12, 20 in arbitrary units. This yields an NTC of 26. With a large total population of *N* = 1000, relative to the NTC, the measured participation ratio of the simulated manifold is 26, and indeed saturates the dimensionality upper bound set by the NTC as predicted.

**Supplementary Figure 10:**
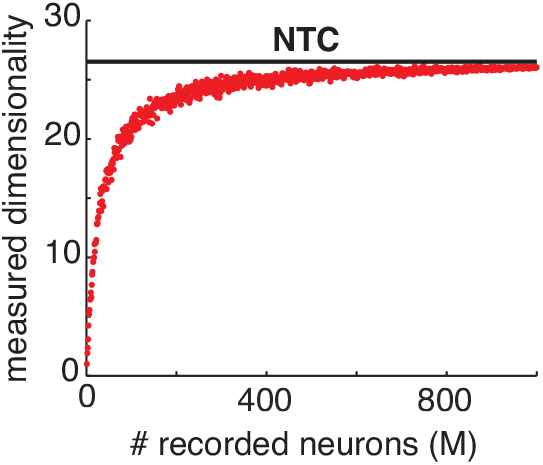
Measured dimensionality of subsampled a random smooth manifold. A random smooth manifold with intrinsic dimensionality of 2 is generated with 1,000 simulated neurons as a Gaussian process with an NTC of 26 (*L*^1,2^ = 100, *λ*^1,2^ = 12, 20). For each value of *M*, a random subset of the simulated neurons is kept to simulated recording. Measured participation ratios of the resulting subsampled data are plotted as function of *M*, and compared against the NTC (horizontal line).

In practice, however, we never record from the entire population of *N* neurons. Suppose we only record a subset of *M* neurons. How would the measured dimensionality scale with *M*, and how large would *M* have to be before the measured dimensionality approached the true dimensionality of the random smooth manifold which is equal to the NTC? We answer these questions by subsampling the number of neurons in the generated random dataset. When plotted as a function of the number of recorded neurons, *M*, (Supplementary Fig. 10), the measured participation ratio of the subsampled data stayed close to the true dimensionality as long as the number of recorded neurons is much higher than the NTC. Only when the number of recorded neurons gets closer to the NTC, does the measured dimensionality start to decrease significantly due to the more stringent constraint of the limited number of recorded neurons.

These results are completely parallel to the case of random clouds of *P* points discussed in the previous section, as opposed to smooth manifolds discussed here. There we found that measured dimensionality monotonically increased with number of recorded neurons *M* in an analytically controllable manner (See Eq. (26)) ultimately saturating as *M ≫ NTC* = *P*. In fact, the red curve in Fig. 10 reflecting the measured dimensionality as a function of number of recorded neurons, is the analog for smooth random manifolds of Eq. (26) for random point clouds.

#### IV.III Intrinsic constraints on neural dynamics can prevent dimensionality from approaching the NTC

In this section, we demonstrate an example simulated dataset that does not saturate the NTC dimensionality upper bound like the *random* smooth manifold. The key idea is to generate neural firing rate patterns that are fundamentally constrained by neural network connectivity. In particular, we build an *N*-dimensional linear dynamical system that evolves according the difference equation,

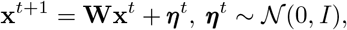

where ***η***^*t*^ is a time-dependent driving input. The length-*N* vector **x**^*t*^ denotes the population activity pattern at time *t*, and the *N*-by-*N* connectivity matrix **W** defines the connectivity of the network. For simplicity, we assume the inputs to be white noise.

**Supplementary Figure 11:**
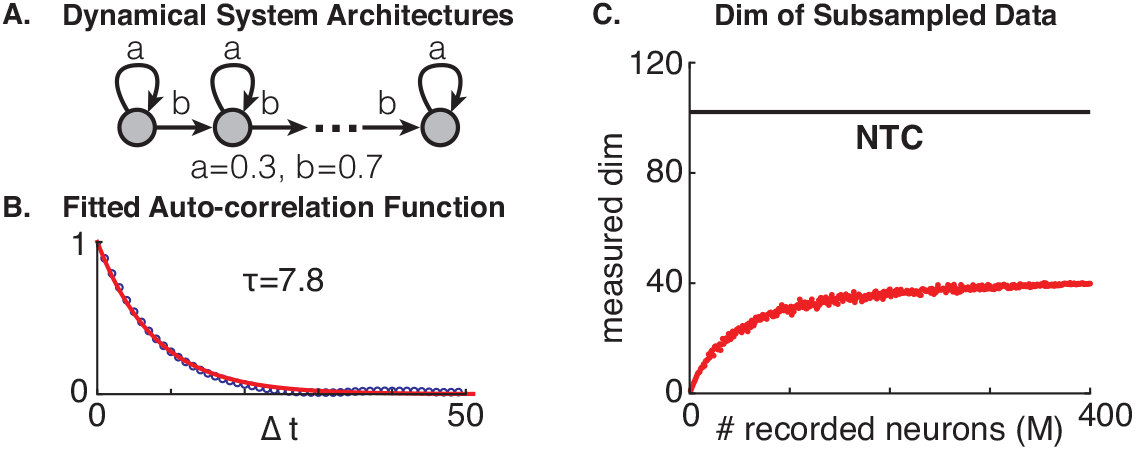
**A**. Architectures of a non-normal linear dynamical system with non-smooth trajectories but low dimensionality. **B**. Fitted auto-correlation function of the simulated data with an exponential function. The data is simulated with *a* = 0.3, *b* = 0.7, *N* = 400 and a duration of 800. **C**. Measured dimensionality of subsampled simulated data compared against the NTC.

We construct a non-normal system with a distributed delay-line architecture (Supplementary Fig. 11A). Its connectivity has the general form,

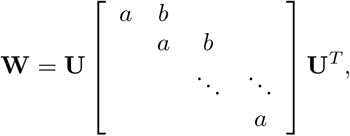

where *a* and *b* are the feedback and feedforward gains at each node; and **U** is an arbitrary orthogonal matrix that distributes the nodes across neurons. With a reasonably large feedforward gain, *b*, input evoked network activity patterns propagate rapidly from one pattern to another associated with successive nodes, resulting in fast-changing single-neuron time courses with a low smoothness value and a high NTC. Despite the non-smooth responses, the system, however, can still be low-dimensional. By the time an input signal propagates to the end of the delay line, it would have incorporated feedforward amplification from all preceding nodes so that the final variance of neural activity along the pattern associated with the last node dominates the total variance of the entire network, resulting in very low dimensionality of the recorded data.

To demonstrate a numerical example, we constructed such a non-normal system with *N* = 400, *a* = 0.3, *b* = 0.7 and a random orthogonal matrix **U**, and simulated recordings of neural activity patterns over a duration of 800 time steps. By construction, the simulated neural activity patterns have a short characteristic time scale of *τ* = 7.8 time-steps, obtained by fitting an exponential auto-correlation function to the stationary approximation of the temporal correlation matrix. The corresponding NTC of the simulated data is then 800*/*7.8 = 102, which is not saturated by the data’s dimensionality of 40, measured by the participation ratio (Supplementary Fig. 11C).

Overall, this model constitutes an example of the elusive experimental regime (iii) in Fig. 4A of the main paper. One need not record all *N* = 800 neurons in the circuit for the dimensionality to stabilize. In fact the dimensionality stabilizes already at *M* = 400 neurons, which is about 10 times the actual dimensionality of 40. So basically, the actual dimensionality is both much less than the number of recorded neurons *and* much less than the NTC - the key signatures of experimental regime (iii). In this case, the gap between the actual dimensionality and the NTC reflects an additional circuit constraint that does not arise from temporal smoothness alone. This constraint corresponds to preferential amplification of a subset of activity patterns associated with the nodes near the end of the hidden delay line.

### V Neural task complexity and random projections as a theory of neural measurement

#### V.I Review of random projection theory

Dimensionality reduction techniques such as principal component analysis attempt to reduce the dimensionality of neural data in a targeted way by computing some data-dependent statistics first, such as the covariance matrix. Alternatively, a dataset may also be dimensionally reduced by simply projecting full data points in a high *N*-dimensional space (in our application with think of *N* as the *total* number of behaviorally relevant neurons in a circuit) into a randomly chosen *M*-dimensional subspace (*M* will correspond to the number of recorded neurons and is much less than *N*). Despite its simplicity, random projections have been shown to nicely preserve the geometric structure of the original high-dimensional data. This structural preservation is measured through the fractional distortion in distance between pairs of data points before and after the projection. Formally, for a pair of *N*-dimensional data points, **x**_*t*_1__ and **x**_*t*_2__ and an *M*-by-*N* random orthogonal projection operator **P**, the pairwise distance distortion is defined as,

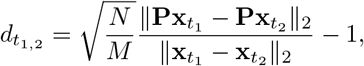

where the ratio 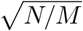 corrects for the expected shrinking of distances under a projection from *N* to *M* dimensions. If this distortion were equal to 0 for all possible pairs, the geometric structure of the dataset would be preserved, since all pairs of points would have the same distance (up to an overall scale) before and after the projection. However, when *M < N*, this is impossible for all pairs of points.

The Johnson-Lindenstrauss lemma [Johnson and Lindenstrauss, 1984, Dasgupta and Gupta, 2003], a fundamental result in the theory of random projections, considers high-dimensional datasets with *P* arbitrary points in the high *N*-dimensional space. The lemma states that, with a probability of at least 1 − *ρ*, a random projection of the *P* data points into a *M*-dimensional subspace has bounded pairwise distance distortions for *all* possible pairs,

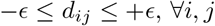

as long as the target subspace’s dimensionality, *M*, is of the order,

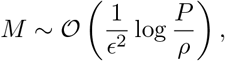

where *ϵ* denotes the maximal, or worse case distortion in distances between all pairs of data points. In other words, the JL lemma tells us that, for random projections to preserve the geometry of a dataset up to some level of tolerance *ϵ* in the fractional error of pairwise distances, it is sufficient to scale the dimensionality *M* of the projected subspace as 1*/ϵ*^2^ and only as the *logarithm* of the number of data points, *P*.

While proven initially for sets of arbitrary data points, the JL lemma has also been extended to datasets with known prior structures. One such extension considers points occupying a *D*-dimensional linear sub-space of the full *N*-dimensional space [Indyk and Motwani, 1998]. The corresponding condition to preserve the geometry of a dataset up to a maximal distortion level *ϵ* requires the projected subspace dimension *M* to scale as

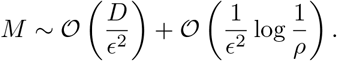

Note that a subspace of dimension *D* is extremely “large”, in the sense that if one wishes to fill a ball of any given radius within the subspace with a cloud of points at a fixed density, the requisite number of points would have to scale *exponentially* with *D*. This phenomenon is one manifestation of the curse of dimensionality; adequate exploration through sampling of a space of dimension *D*, without missing any regions, requires a number of samples that grows exponentially with *D*. However, despite this curse of dimensionality, the number of random projections *M* required to preserve the geometry of *all* pairs of points within a subspace of dimension *D* at some fixed distortion *ϵ*, need only scale *linearly* with *D*. Roughly, the subspace JL lemma follows from the pointwise JL lemma by making the replacements *P → e*^*D*^ and log *P → D*. In this sense they are consistent with each other.

A similar extension considers data points sampled from a low *K*-dimensional non-linear manifold with a finite volume *V* embedded in an *N*-dimensional space [Baraniuk and Wakin, 2007]. The sufficient condition to preserve the geometric structure of the data to within fractional distortion *ϵ* requires the subspace dimension *M* to scale as,

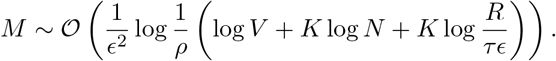

This scaling is similar to the linear subspace case, as it grows logarithmically with the volume of the manifold. Given that the volume of a manifold grows exponentially with the intrinsic dimension (log *V ∝ K*), the scaling of *M* is, in fact, linear in the dataset’s intrinsic dimension *K*. Different from the previous two cases, however, the random projection of manifolds includes an addition dependency on the embedding dimensionality, *N*, which we will check in the next section with simulations. The condition number *τ* and geodesic covering regularity *R* characterizes the curvature of the nonlinear manifold, which are properties of the manifold’s geometry.

These theoretical results all suggest that random projections yield extremely efficient representations of high-dimensional data, as long as the number of coefficients in the reduced representation grows logarithmically with the amount of data. With different theorems, this amount of data is measured in different ways: by the number of data points in the JL lemma, by the exponentiated subspace dimension for linear subspaces, and by the volume of a nonlinear manifold.

In the main paper, we argue that, in many scenarios, random projection theory constitutes a good model of the process of neural measurement of a random subset of *M* neurons. Conceptually, the process of projecting high-dimensional data points into a randomly generated low-dimensional subspace in a random and data-blind way is similar to the random sampling of the high-dimensional patterns of neural activity across all *N* neurons in a circuit, down to a random subset of *M* neurons that happen to be measured in a single experiment. Indeed, when the full *N*-dimensional neural activity patterns are randomly oriented with respect to the single neuron axes, the neuroscientists’ act of random sampling is exactly equivalent to the mathematical procedure of a random projection.

To see this equivalence, consider a full-dimensional dataset denoted as the *N*-by-*N*^*T*^ data matrix **X**. If we force the data to be randomly oriented by applying a random orthogonal rotation to obtain a rotated dataset, **UX**, the random sampling of its rows is mathematically equivalent to the random projection of the original data with the sampled rows of **U** as the projection’s basis. Consequently, the attractive scaling laws governing the requisite number of random projections *M* to achieve a desired distortion *ϵ* can be directly translated into scaling laws governing the required number of neurons to record in order to achieve a given desired accuracy of the resultant dynamic portraits of circuit computation obtained via dimensionality reduction. Of course, this equivalence comes with the caveat that neural activity patterns should be randomly oriented with respect to the single neuron axes. In such a scenario, neural activity patterns are distributed, or exhibit mixed selectivity in which many neurons code for many task parameters. A major departure from this assumption is a high degree of sparsity, and we will investigate the effects of sparsity using simulations below.

#### V.II Numerical analysis of scaling behavior a random projection theory of measurement

Since, as nonlinear representations of stimulus or behaviorial variables, trial-averaged neural data can be best described as nonlinear manifolds embedded in the full *N*-dimensional firing rate space of all neurons in the circuit, we verify the scaling laws of the random projection of smooth manifolds in this section using simulations. We further aim to make the proportionality constants concrete in the sufficient condition for accurate recovery,

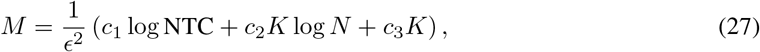

where we replaced the manifold volume, *V*, with NTC, since they are scaled versions of each other. Because NTC scales exponentially with the intrinsic dimensionality, *K*, this sufficient target dimension *M* is still linearly proportional to *K*.

We first check the scaling against log NTC using simulated data from one- and two-dimensional smooth manifolds generated using Gaussian processes. With the kernel’s characteristic time scale fixed at, *τ* = 12, the NTCs for the 1D and the 2D cases are simply 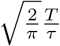 and 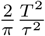 (the 2D Gaussian process has a factored kernel), where *T* is the range of the task parameters. In both cases, we embedded the generated random manifold in a 1,000-dimensional space, and computed the maximal pairwise distortions, 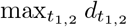, for all possible pairwise distances under random projections of different *M*s and of different NTCs (by varying *T* from 4 to 100). For each combinations of *M* and NTC, we computed the maximal distortion over 100 trials, and used the 95 percentile (*ρ* = 0.05) of all the maximal distortions as the value for *ϵ*.

With the distortion, *ϵ*, plotted in the *M*-vs-log NTC plane, we highlighted values of *M* for three different distortion levels, *ϵ* = 0.2, 0.3, 0.4, at different NTCs, and obtained excellent linear fits of the corresponding constant-distortion contours as predicted (Supplementary Fig. 12A). Furthermore, with a renormalization of the highlighted *M* values by *K/ϵ*^2^, the fitted linear functions of log NTC*/K* under different parameter combinations have similar slopes, or *c*_1_ (Supplementary Fig. 12B). Indeed, the slopes fitted using 1D manifolds have an average of 1.07 ± 0.07, which is within the region of uncertainty for the average slopes fitted using 2D manifolds, 1.05 ± 0.11.

**Supplementary Figure 12:**
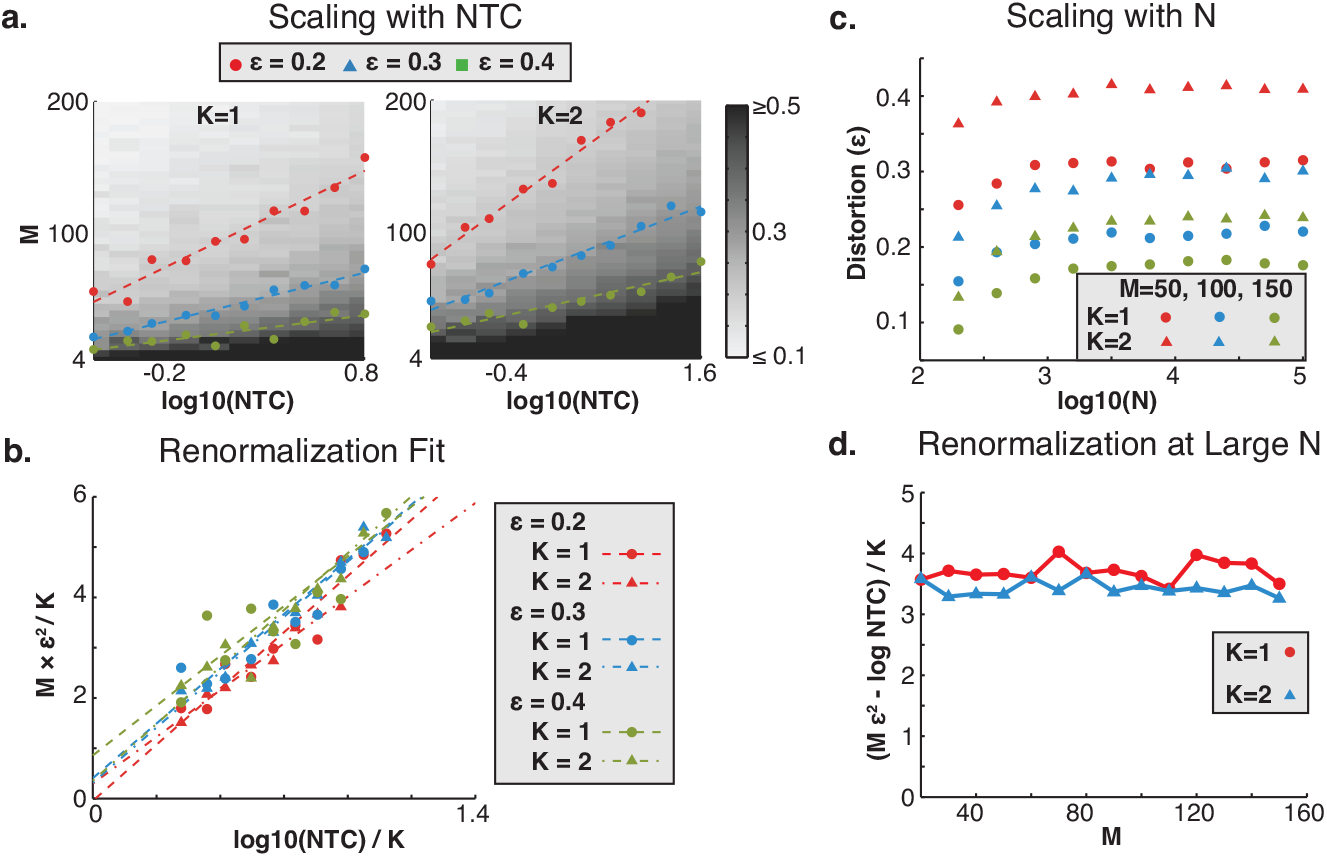
Random projections of simulated smooth manifold. **A**. Simulated distortions, *ϵ* with *ρ* = 0.05, of smooth random manifolds generated using 1D and 2D (factored kernel) Gaussian processes. Distortions are plotted in the plane of the number of recorded neuron, *M*, versus the logarithm of the NTC. Locations of exemplar distortions, *ϵ* = 0.2, 0.3, 0.4 are highlighted and fitted with linear functions of log NTC. **B**. Fitted constant-distortion contours on rescaled x and y axes to extract the parameter *c*_1_. **C**. Distortion’s saturation as a function of the embedding dimension, *N*, for 1D and 2D manifolds with *M* = 50, 100, 150. **D**. Offset and rescaled distortions as a function of *M* in the limit of large *N* = 100, 000.

Next, we check the scaling of *M* with the dimension of the embedding space, *N*. Since simulations become very expensive for high embedding dimensions, we compute the distortions under just three values of *M* = 50, 100, 150 for embedding dimensions ranging logarithmically from 200 to 100,000 (Supplementary Fig. 12C). While the simulated distortions seem to increase linearly with log *N* initially, as the embedding dimension becomes large (*N* ~ 10, 000), the distortion *ϵ* derived from simulations saturate to a constant in each case. While seemingly contradictory, the theory isn’t falsified since it provides only a *sufficient* condition on the scaling of *M*, and would be consistent with any scaling that is better than log *N* (Eq 27).

So what happens at large *N*? Can we tighten the published sufficient condition to better describe our simulations of the random projection of smooth manifolds? Given our observation of the saturating behavior, a simple modification of the scaling law is to simple replace the log *N* term with a constant, which we combine with *c*_3_ to obtain the following simplified scaling law,

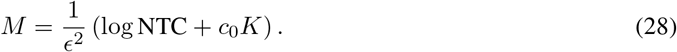

In this new formula, we have also replaced the *c*_1_ term with our fitted value of 1. Given this hypothetical relation, the constant *c*_0_ must then equal to 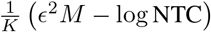 for large *N*. To check its validity, we simulated additional 1D and 2D manifolds in 100,000-dimensional embedding spaces with values of *M* varies more densely from 10 to 150 (Supplementary Fig. 12D). Remarkably, for combinations of different *M* and *K* values, the resulting values of *c*_0_ remain extremely steady around a value of 3.5 ± 0.2, consistent with our simplified scaling law.

#### V.III Scaling of distortion with neural sparsity

While random projection and random sampling are mathematically equivalent when the neural activities are randomly oriented with respect to the neuronal axes, actual neural data may be more sparse and preferentially oriented along the neuronal axes. To quantify how distributed or sparse a set of *M*-dimensional population activities are, we use a measure called data coherence, which, for the *M*-by-*N*^*T*^ data matrix **X**, is defined as,

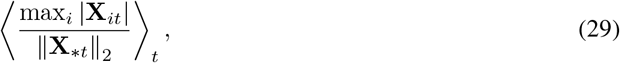

where **X**_*∗t*_ denotes the *t*th column vector of the data matrix. Algebraically, the data coherence measure averages the individual column vectors’ sparsity, which is defined as the ratio of the vector’s coordinate with the maximal magnitude to its norm. Conceptually, the data coherence measure quantifies a dataset’s sparsity by computing its mutual coherence–a concept widely utilized in the compressive sensing literature–with respect to the neuronal basis. Numerically, this measure of sparsity varies from 1 for a perfectly sparse dataset, where only a single neuron is active in any recorded population activity pattern, to 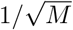 for a distributed dataset, where components of the population activity are evenly distributed across neurons. The chance level coherence of a zero mean unit variance random gaussian dataset is roughly 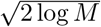, which is the expected maximum for the absolute values of *M* zero mean unit-variance gaussian random variables. Note that for the same underlying network size, *N*, the data coherence measure may change depending on the number of recorded neurons.

To investigate how sparsity affects the number of required neurons for the accurate recovery of dynamical portraits under subsampling, we use simulations of nonlinear neural networks whose activities patterns’ sparsity can be controlled. The simulated recurrent nonlinear neural networks evolve according to the following differential equation,

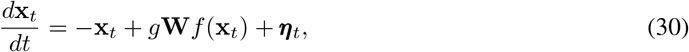

where the state of the *N* neurons in the network at time *t* is represented by the vector **x**_*t*_. Neuronal dynamics is modeled using the soft-thresholding nonlinearity,

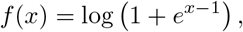

and produces outputs which are then fed through the connectivity matrix **W** with some gain factor *g* into the rest of the network. The network is driven by random *N*-dimensional pink noise, ***η***_*t*_, smoothed with a gaussian kernel of a width of 5ms. For the network dynamics to dominate the neuronal dynamics, we fix *g* at 0.99 for our simulations. To control the sparsity of the simulated activities, we manipulate the sparsity of the connectivity matrix using the following formula,

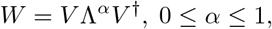

where the product of *V* Λ*V*^†^ is the eigen-decomposition of a randomly generated orthogonal matrix obtained from applying the Gram-Schmidt procedure to a random gaussian matrix. The parameter *α* corresponds to an element-wise exponentiation of the decomposition’s eigenvalues, which are pure imaginary numbers.

As *α* varies from 0 to 1, *W* transitions smoothly from the sparse identity matrix to a distributed random orthogonal matrix, generating neural activity patterns of varying sparsity.

**Supplementary Figure 13:**
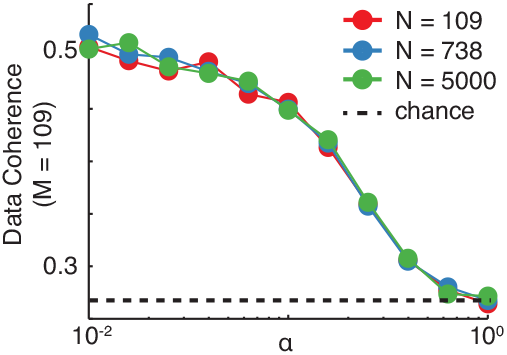
Relation between the parameter *α* and the resulting data coherence measured with a subset of 109 neurons for different network sizes.

We empirically mapped the relation between the parameter *α* to the simulated activities’ data coherence with networks of different sizes 109, 738, and 5000 run with values *α* varied logarithmically from 0.01 to 1 (Supplementary Fig. 13). When measured with a fixed 109 number of neurons (same as monkey H), the measured coherences of samples activities depended only on the parameter *α* but not the network size, suggesting that modifying the value of *α* is a controlled way of exploring data coherence without affecting other properties of the simulated neural activity patterns. For the rest of this section, we compare random samplings of simulated activities with coherences of either 0.27 or 0.55 inclusive of observed coherence in data (0.34 for monkey H and 0.43 for monkey G).

Applying both random samplings and random projections to the simulated dataset, we explored their resulting distortions in the *M*-by-log *T* plane, and fitted to them constant distortion contours of the same three levels (Figure 4B,C). The excellent linear fits of *M* against log *T* across data coherences for both random projections and random sampling reaffirms that the logarithmic scaling of the requisite number of neurons is a robust phenomenon that persists across a large range of sparsity levels of neural activity patterns.

Furthermore, comparing results in the case where simulated neural activity patterns are randomly distributed (data coherence = 0.27, leftmost panels, Figure 4A,B,C,D), we see no difference between the requisite number of neurons for random sampling and the requisite subspace dimension for random projection, verifying the intuitive connection we’ve made between the two for randomly oriented activity patterns. The increase in the number of requisite neurons relative to the number of random projections only starts to appear as simulated neural activity patterns become more sparse, in agreement with the small discrepancy observed in the motor cortical data (Figure 3H) where neural activities have less-than-random orientations with respect to the neuronal axes as reflected by their higher data coherence (0.32 *>* 0.27).

More quantitatively, we compared, for the distortion value of 0.3, the slopes (increase in the requisite number of neurons or random subspace dimension for every 10-fold increase in volume) and intercepts (the minimal number of neuron or subspace dimension) of the *M* versus log *T* fits as functions of data coherence for random sampling and random projection. We see that the slopes and intercepts in the case of random sampling increased systematically with data coherence, or sparsity, reflecting the increased chance of missing “important” neurons as trajectories become sparser (Figures 3A,B). However, at data coherence levels around the value observed in our motor cortical dataset, the increases in both the fitted slope and intercept are quite small, suggesting that the motor cortical data are not far from being randomly oriented. On the other hand, because random projection always results in subspaces that are, by definition, randomly oriented with respect to any chosen basis of the firing-rate space, neither the fitted slopes nor the intercepts have a dependency on the simulated activities’ data coherence. This behavior is consistent with the fact that properties other than sparsity are well controlled in our simulations.

**Figure 14:**
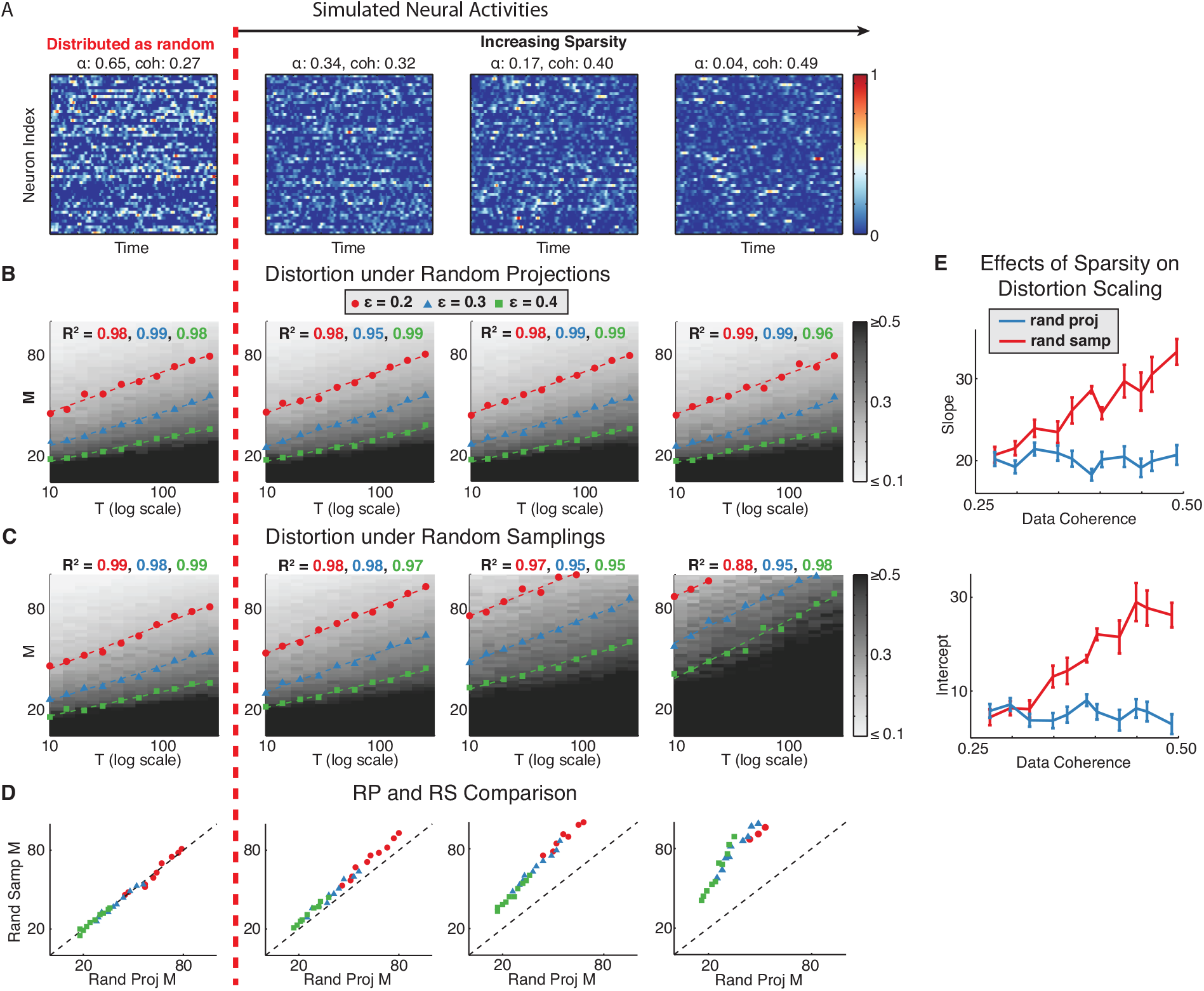
Effect of sparsity A. Example firing rates of simulated networks with different levels of sparsity; **B**. Constant distortion contours for the simulated activities under random projection; **C**. Constant distortion contours for simulated activities under random sampling; **D**. Comparisons of the requisite number of neurons under random projections and random samplings. **E**. Fitted slopes and intercepts of constant distortion contours as functions of data coherence. Error bars correspond to standard errors of the fitted parameters.

Overall, analysis of simulated data suggests that the optimistic result that the number of recorded neurons need only scale logarthmically with the neural task complexity to achieve a fixed distortion in neural state space dynamics, holds true for a broad range of sparsities of neural activity patterns. The actual requisite number of neurons grows with increasing sparsity, but the linear relation between *M* and log NTC is preserved. While we support this conclusion using simulation results, it is a highly plausible theoretical conjecture as well. In fact, recent theoretical progress has been made to extend the basic JL lemma beyond random projections into cases where the projection vectors are sparse [Achlioptas, 2003, Li et al., 2006], including random sampling.

## Appendix

### Analytical expressions for fraction of variance explained

For the exponential correlation function, 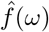 evaluates to,

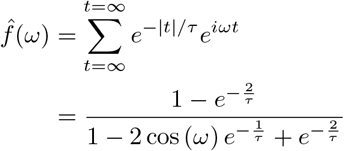

Taking advantage of 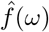’s ordering, we evaluate the integral,

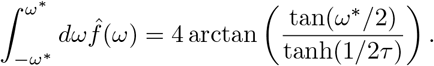

The minimum *ω\** given the fraction parameter *r* is then,

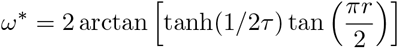

With the *τ* ≫ 1, we ignore terms beyond 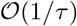 to obtain the clean, final expression,

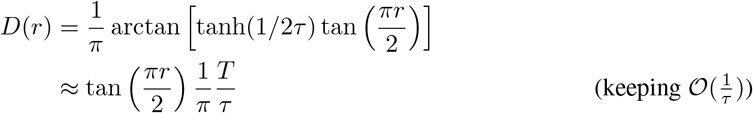

For the gaussian temporal correlation function, we state without showing that the Fourier transform is in fact nicely ordered. To compute the fraction of variance explained, we switch the order of the summation over *t* and the integration of *ω* to obtain,

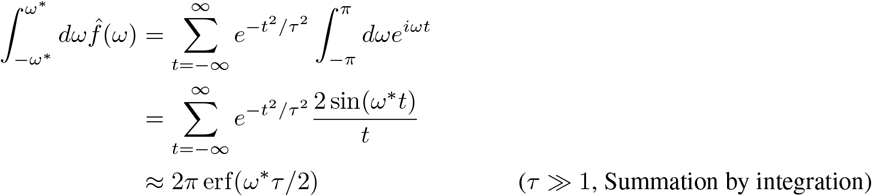

With the optimal *ω\** ≈ 2 inverf(*r*)*/τ*, we then have the final expression,

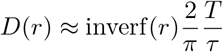

